# Subventricular zone/white matter microglia reconstitute the empty adult microglial niche in a dynamic wave

**DOI:** 10.1101/2021.02.17.431594

**Authors:** Lindsay A. Hohsfield, Allison R. Najafi, Yasamine Ghorbanian, Neelakshi Soni, Joshua D. Crapser, Dario X. Figueroa Velez, Shan Jiang, Sarah E. Royer, Sung Jin Kim, Aileen J. Anderson, Sunil P. Gandhi, Ali Mortazavi, Matthew A. Inlay, Kim N. Green

## Abstract

Microglia, the brain’s resident myeloid cells, play central roles in brain defense, homeostasis, and disease. Using sustained colony-stimulating factor 1 receptor inhibition, we report an unprecedented level of microglial depletion and establish a model system that achieves an empty microglial niche in the adult brain. We identify a myeloid cell that migrates from an important neurogenic niche, the subventricular zone, and associated white matter areas. These cells exhibit tremendous chemotaxis potential, migrating radially and tangentially in a dynamic wave and filling the brain in a distinct pattern, to fully replace the microglial-depleted brain. These repopulating cells are enriched in disease-associated microglia genes and exhibit distinct phenotypic and functional profiles to endogenous microglia. Our findings shed light on the overlapping and distinct functional complexity and diversity of myeloid cells of the CNS and provide new insight into myeloid cell dynamics in an empty microglial niche without contributions from bone marrow-derived cells.

## Introduction

Microglia represent the largest population of immune cells in the brain, constituting 5-10% of brain cells in the adult central nervous system (CNS). As resident tissue macrophages, microglia are responsible for immune defense and resolution, tissue maintenance, neuronal support, and synaptic integrity^1–3^. Their central role in the CNS makes microglia attractive drug targets for neurological disorders/injuries. However, developing effective therapies that manipulate microglia requires further understanding of microglial origins, diversity, homeostasis, and dynamics.

Microglia arise from yolk sac-derived erythromyeloid progenitors and colonize the brain as embryonic microglia during early stages of development (i.e. E8.5 – E9.5)^4, 5^. These immature myeloid cells, displaying amoeboid morphology and high proliferative potential, enter the brain via the meninges and ventricles in mice^6–9^, as well as, via the leptomeninges, choroid plexus, and ventricular zone in humans^2, 10–12^. Microglial colonization occurs first in the white matter (WM) (e.g. internal capsule, external capsule, and cerebral peduncle) and continues to the sub- and then cortical plate as cells proliferate and migrate in a radial and tangential manner^10^. In adulthood (P28 and onward), microglia become fully mature, exhibiting ramified morphology and expressing canonical microglial signature genes: *P2ry12*, *Tmem119*, *Siglec-H*, *Cx3cr1*, *Olfml3*, *Fcrls*, and *Sall1*^13, 14^.

Recent single-cell RNA sequencing studies have identified transcriptionally distinct microglial gene signatures associated with disease (e.g. disease-associated microglia (DAM), microglial neurodegenerative (MGnD) phenotype^15–18^), injury (e.g. injury-responsive microglia (IRM)^19^), and brain region-specific areas/developmental stages (e.g. proliferative-region-associated microglia (PAM), axon tract-associated microglia (ATM)^18–, 21^). Despite this, homeostatic microglia appear less heterogeneous during adulthood^19, 20^. These findings shed light on microglial diversity and state changes during health and disease, however, it remains unclear whether adult homeostatic microglia exist as one population, with the ability to change from one transcriptional/functional state to another, or whether they exist as heterogeneous subpopulations with distinct propensities.

Microglial homeostasis and dynamics are maintained by many signaling factors, including transforming growth factor-beta, Il-34, and colony-stimulating factor 1^22, 23^. Recent studies exploring the homeostatic kinetics of the microglia in the adult brain have revealed that these cells are long-lived^2, 24^ and self-renew, even after acute 80-95% depletion and subsequent repopulation^23, 25, 26^. However, no approach to date has been able to deplete all microglia^27, 28^, and the rapid proliferation of surviving microglial cells would obscure the detection of other myeloid cells that contribute to the CNS environment and/or repopulation in the adult brain. While conventional depletion paradigms have shown that microglia have a remarkable capacity to repopulate from the presence of few surviving cells, we sought to investigate the consequences of eliminating these few remaining cells on microglial population dynamics.

To address this, we have optimized a colony-stimulating factor 1 receptor (CSF1R) inhibitor approach that involves sustained inhibitor administration, building on our prior work that microglia are dependent on this signaling for their survival^23^. This approach results in a delayed repopulation of myeloid cells that reconstitute the brain in a sequential manner previously unseen in the adult brain. We show that repopulating cells emerge from the subventricular zone (SVZ) and traffic throughout the brain parenchyma via WM tracts before spreading out radially and tangentially through the rest of the brain in a dynamic wave of proliferating cells. Following full brain reconstitution, these repopulating cells remain phenotypically, transcriptionally, and functionally distinct from endogenous microglia, demonstrating unique gene expression profiles that are enriched for DAM genes and a unique capacity to significantly reduce amyloid burden in a mouse model of AD. Together, these data identify a previously undescribed novel source of brain myeloid cell repopulation - highlighting a distinct repopulation paradigm that provides further insight into study of myeloid cell homeostasis, dynamics, and disease.

## Results

### Sustained high dose of CSF1R inhibitor unmasks a distinct form of myeloid cell CNS repopulation

In previous studies, we have shown that 7d treatment of the brain penetrant CSF1R/KIT/FLT3 inhibitor PLX3397 (Pexidartinib; 600 ppm in chow) eliminates ∼90-98% of microglia in the CNS^23, 29^. During depletion, surviving microglia are seen scattered throughout the brain (Figure 1A-B, C) and subsequent withdrawal of the inhibitor results in rapid and spatially homogenous microglial repopulation within 3d, with cells exceeding control numbers by 7d (Figure 1C, E). Recent studies show that repopulation is dependent on the local proliferation and clonal expansion of surviving microglia^23, 25, 26, 30^; thus, we refer to this type of repopulation as partial microglial niche-dependent (PND) repopulation.

**Figure 1.**
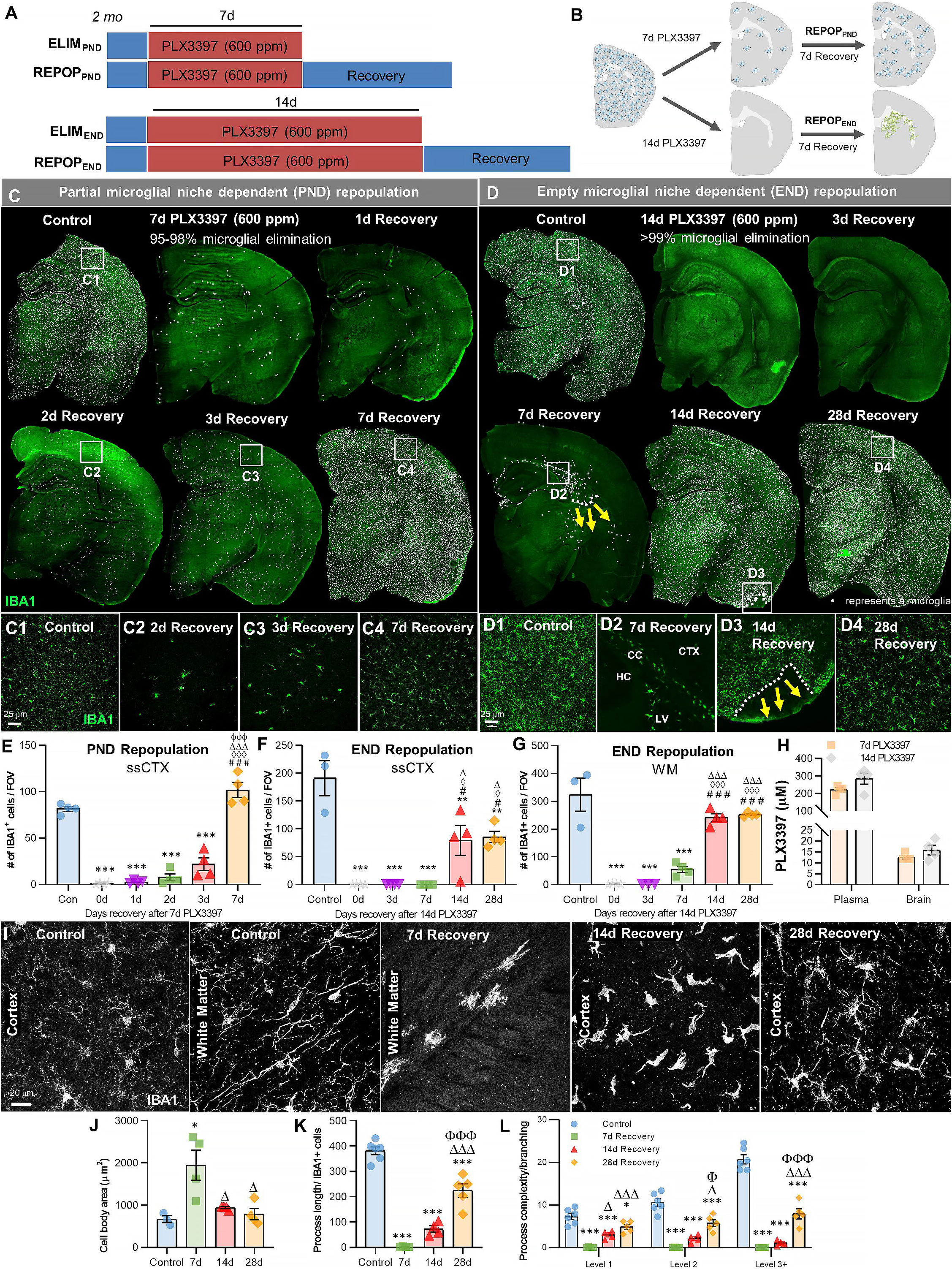
**Sustained high dose of CSF1R inhibitor unmasks a distinct form of myeloid cell CNS repopulation.** (A-B) Experimental paradigm and schematic depicting dose and duration of PLX3397 (600 ppm) treatment and subsequent inhibitor withdrawal allowing for partial niche-dependent (PND) and empty niche-dependent (END) microglial repopulation. For PND repopulation: 2-month-old wild-type (WT) mice were treated with 600 ppm of PLX3397 for 7d, achieving ∼90-98% brain-wide microglial depletion, with remaining microglia visibility dispersed throughout the brain parenchyma, and then placed on control diet for 7d (7d recovery) allowing for microglial repopulation. At 7d recovery, repopulating microglia reconstitute the brain from areas in which previously remaining microglia were deposited. For END repopulation: two-month-old WT mice were treated with 600 ppm of PLX3397 for 14d, achieving 99.98% brain-wide microglial depletion, and then placed on control diet for 7d allowing for microglial repopulation. At 7d recovery, repopulating myeloid cells reconstitute the brain in specific neuroanatomical niches (e.g. ventricle, subventricular zone, white matter tracts, caudoputamen). (C-D) Representative immunofluorescence whole brain images of myeloid cells (IBA1, green) at each time point of treatment and recovery during PND (C) and END (D) repopulation, with white dots superimposed over microglia. Due to the differential kinetics of these two forms of repopulation, repopulation (i.e. recovery) was evaluated at different timepoints, including initial stages with few cells, mid-repopulation, and full brain reconstitution. For PND repopulation: 2-month-old WT mice treated with control, 7d PLX3397, 7d PLX3397 followed by 1 day on control diet (1d Recovery), 7d PLX3397 followed by 2 days on control diet (2d Recovery), 7d PLX3397 followed by 3 days on control diet (3d Recovery), and 7d PLX3397 followed by 7 days on control diet (7d Recovery). For END repopulation: 2-month-old WT mice treated with control, 14d PLX3397, 14d PLX3397 followed by 3 day on control diet (3d Recovery), 14d PLX3397 followed by 7 days on control diet (7d Recovery), 14d PLX3397 followed by 14d days on control diet (14d Recovery), and 14d PLX3397 followed by 28 days on control diet (28d Recovery). (C1-C4, D1-D4) Inserts of higher resolution confocal images of IBA1^+^ cells during repopulation. White dotted lines and yellow arrows highlight “wave” edge and direction. (E-G) Quantification of IBA1^+^ cells per field of view (FOV) at each time point in cortical and white matter regions, respectively during PND (E) and END (F-G) repopulation. (H) Pharmacokinetics analysis of PLX3397 levels in plasma and brain of mice treated with 7d and 14d PLX3397 (600 ppm). (I) Representative 63x immunofluorescence images of myeloid cells (IBA1, white) display morphological alterations. (J-L) Quantification of IBA1^+^ cell morphology: cell body area in the white matter tract (J), average process/filament length (K), and process complexity/branching (L) in the piriform cortex. Level 1-3+ indicates level of branching from the cell body. Data are represented as mean ± SEM (n=3-4). *p < 0.05, ** p < 0.01, *** p < 0.001; significance symbols represent comparisons between groups: (E) control *, 0d #, 1d ◊, 2d Δ, 3d Φ; (F-G) control *, 0d #, 3d ◊, 7d Δ. CC, corpus callosum; CTX, cortex; HC, hippocampus; LV, lateral ventricle.

Here, we set out to examine CNS myeloid cell repopulation dynamics in the absence of remaining microglia in the brain. To accomplish this, we utilized a sustained high dose of PLX3397 (600 ppm) for 14d. This treatment results in pharmacologically unprecedented microglial depletion, in which we observe no IBA1^+^ cells across whole brain sections (Figure 1A-B, D). Although PLX5622 is a CSF1R inhibitor (CSF1Ri) that is more active against CSF1R compared to other related kinases, studies in our lab have shown that high dose PLX3397 vs high dose PLX5622 achieves higher CNS exposure and microglial depletion efficiency. Pharmacokinetic analysis of microglia-depleted brains at both 7d and 14d treatment of PLX3397 shows that PLX3397 levels remain the same in the CNS despite longer drug exposure (Figure 1H).

To explore the differential repopulation dynamics between these microglial elimination paradigms, we treated mice with PLX3397 for 14d and then withdrew the inhibitor, allowing the CNS to recover for 3d, 7d, 14d, and 28d (Figure 1D, F-G), and compared it to PND repopulation (Figure 1C, E). At 3d recovery following 14d PLX3397 treatment, no IBA1^+^ cells are detectable in most brain sections. By 7d recovery, IBA1^+^ cells appear, but are exclusively located near the lateral ventricle and in WM tracts lining the ventricles (Figure 1D2). By 14d recovery, IBA1^+^ cells have spread throughout most of the CNS; however, some areas of the cortex (e.g. the piriform cortex) remain unoccupied (Figure 1D3). These areas display a distinct “wave” of cells in adjacent unoccupied cortical areas (Figure 1D3). At 28d repopulation, all brain regions are populated with IBA1^+^ cells, but absolute cell numbers remain 50% lower compared to microglia in control animals, as seen in somatosensory cortices (Figure 1D4, F). We subsequently refer to this form of repopulation as empty microglial niche dependent (END) repopulation, due to its distinct characteristics from PND repopulation.

We have previously shown that within 14-21d of PND repopulation, repopulating microglia not only attain similar densities to resident microglia, but also display similar morphologies, cell surface marker expression, gene expression profiles, and inflammatory responses to LPS^23, 31, 32^. In contrast, END repopulating cells display larger cell bodies (Figure 1I-J), reduced process/filament length (Figure 1I, K), and reduced process/dendrite branching and complexity (Figure 1I, L) compared to homeostatic microglia even after 28d recovery.

### Extensive CSF1R inhibition unveils the presence of CSF1Ri-resistant myeloid cell in the subventricular zone/white matter areas

Having demonstrated that 14d PLX3397 (600ppm) treatment and subsequent withdrawal results in reconstitution of the adult brain with a phenotypically distinct myeloid cell with unique tempo-spatial migratory patterns, we next sought to determine the source of these cells. To that end, we first confirmed the extent of microglial depletion with multiple myeloid markers, including microglial-specific P2RY12 and TMEM119 (Figure 2A-B), as well as in *Cx3cr1^CreERT2^* mice, to explore whether surviving cells were present but just downregulating myeloid markers. In these mice, YFP is permanently expressed in microglia following tamoxifen-inducible lineage tracing, illustrating that depletion is not due to a downregulation in microglial markers, but a loss of cells (Figure 2C). While in previous analyses we observed no IBA1^+^ (including Cd11b^+^, P2RY12^+^, and TMEM119^+^) cells throughout the brains following 14d PLX3397 treatment, we next conducted an examination throughout the entire brain along the rostral-caudal axis (i.e. every 6^th^ section). With this extensive analysis, we observe a very small number of surviving IBA1^+^ cells in treated brains (∼15 in 14d PLX3397 brains vs. ∼132,000 in control brains = 99.98% depletion; Figure 2D). Despite this, we report the highest reported loss of microglial cells in the adult brain. Notably, these few cells (0.02% of cells) are seen exclusively in ventricular (i.e. SVZ) and adjacent WM areas (Figure 2E-F). Moreover, these cells displayed a lack of staining for P2RY12 and distinct morphological profiles (Figure 2G-H). Reports show that the population of microglia residing in the adult SVZ and adjacent RMS display a distinct morphological profile with an amoeboid cell body and fewer/shorter branched processes and exhibit an activated phenotype, similar to END repopulating cells^33–35^. Prior descriptions of myeloid cells found in the adult SVZ have found lower expression levels of the microglial-specific marker P2RY12^33^. Here, we find repopulating cells are initially negative for both microglial-specific P2RY12 and TMEM119 surface markers, however, express these markers by 28d recovery (Figure 2I-J, Figure S1A).

**Figure 2.**
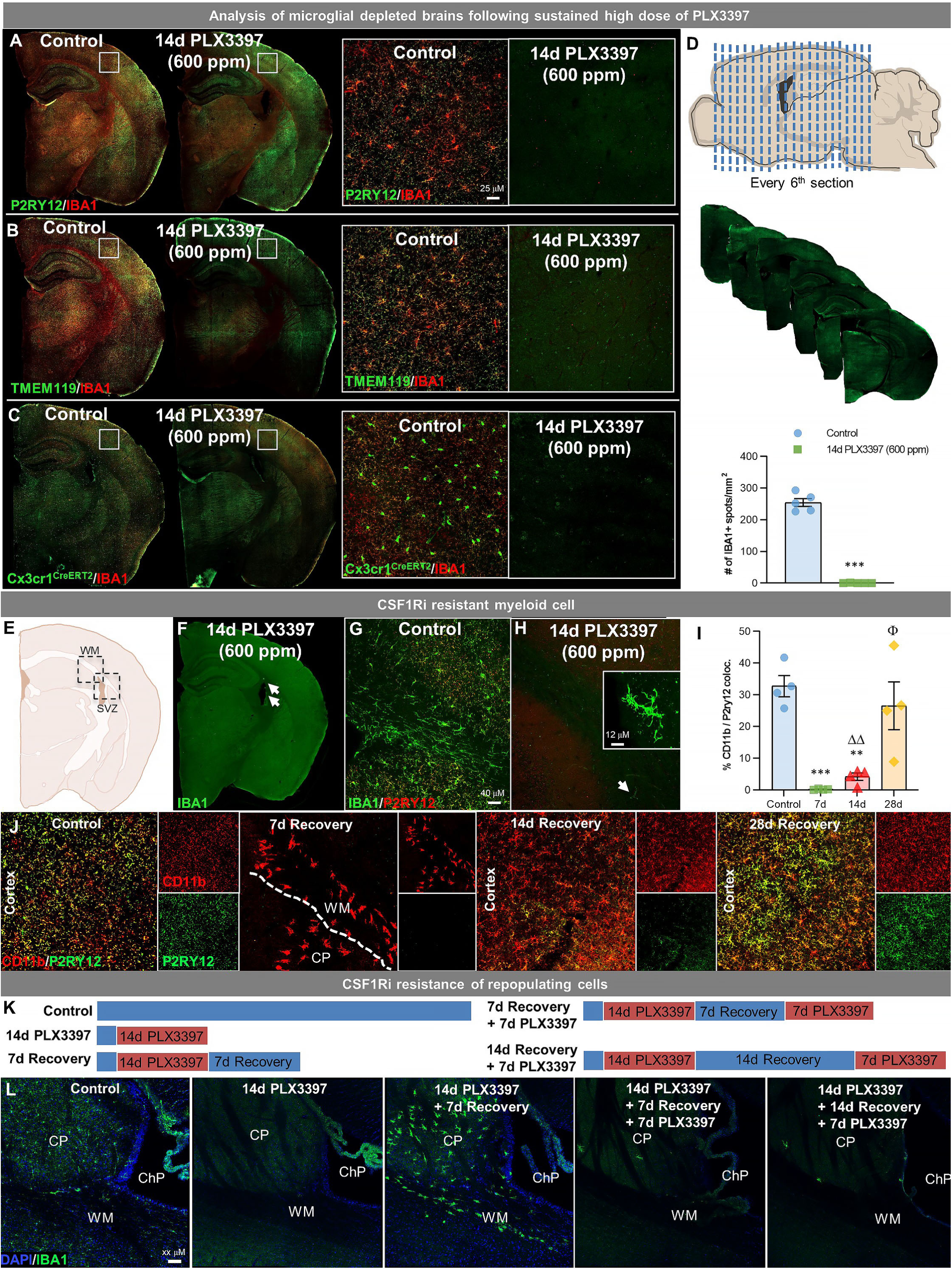
**Extensive CSF1R inhibition unveils the presence of CSF1Ri-resistant myeloid cells in the subventricular zone and white matter tracts.** Two-month-old WT or *Cx3cr1^CreERT2^* mice treated with vehicle or PLX3397 (600 ppm in chow) for 14 days to evaluate extent of microglial depletion. (A-C) Representative whole brain slice images of IBA1^+^ cells co-stained with microglial-specific markers P2RY12 (A) and TMEM119 (B), or YFP, to detect microglia or cells of *Cx3cr1* lineage (C), and ultimately microglial depletion efficiency. For *Cx3cr1* lineage tracing studies, *Cx3cr1^CreERT2^* mice were crossed with YFP reporter mice, treated with tamoxifen and then allowed to recover for 21 days prior to PLX3397 treatment. Inserts show 20x high resolution images of brain regions highlighted by white boxes. (D) Total IBA1^+^ cells were quantified throughout the entire brain taking every 6^th^ section (i.e. stereology-style) along the rostral-caudal axis of 2-month-old mice treated for 14d with PLX3397 (600ppm in chow). (E) Brain section schematic of SVZ and WM areas where CSF1R-resistant myeloid cells are present following 14d PLX3397 (600ppm in chow) treatment. (F) Representative whole brain slice image of CSF1R-resistant IBA1^+^ cells (white arrows) near the SVZ/WM areas surrounding the lateral ventricle. (G-H) Representative 20x immunofluorescence image of IBA1^+^ cells co-stained with P2ry12 in control (G) and 14d PLX3397 (H) mice, showing that CSF1R-resistant cells are P2RY12^-^. Insert shows higher resolution image of CSF1R-resistant myeloid cell illustrating their atypical morphological profile. (I) Quantification of % colocalization of CD11b^+^ and P2RY12^+^ cells as seen in (J). (J) Representative 20x images of myeloid cells (Cd11b, red) and a microglial-specific marker (P2RY12, green). (K) Experimental schematic depicting dose and duration of CSF1Ri resistance study. Two-month-old WT mice were placed on PLX3397 (600 ppm) for 14d, allowed to recover for 7d or 14d on control diet (allowing for END repopulation), and then placed back on CSF1Ri diet (PLX3397 600 ppm) for 7d. Controls, 14d PLX3397, and 7d Recovery were also included for comparison. (L) Representative immunofluorescent images of IBA1^+^ cell (green) and DAPI staining (blue, used to distinguish structures), showing the susceptibility of END repopulating cells once migrated from SVZ/WM areas. Data are represented as mean ± SEM (n=3-5). *p < 0.05, ** p < 0.01, *** p < 0.001; significance symbols represent comparisons between groups: control *, 0d #, 3d ◊, 7d Δ, 14d Φ. CP, caudoputamen, ChP, choroid plexus, WM, white matter.

To explore whether END repopulating microglia maintain resistance to CSF1Ri following their migration from the SVZ, we treated 7d and 14d recovery mice with 7d PLX3397 (600 ppm) (Figure 2K). At both recovery times, the majority of cells were eliminated (Figure 2L) showing that END repopulated cells are susceptible to CSF1Ri treatment and require CSF1R signaling for their survival.

To conclusively determine whether this unique form of repopulation occurs as a result of the unprecedented level of microglial depletion vs. a 14d requisite CSF1Ri drug treatment, we utilized H2K-BCL2 mice, in which overexpression of BCL2 in myeloid lineages affords some resistance to CSF1Ri-induced cell death. In these mice, 14d treatment with PLX3397 (600 ppm) leads to a 90-95% elimination of microglia (similar to a 7d treatment in WT mice), and subsequent withdrawal of drug elicits PND repopulation, rather than END repopulation (Figure S1B-C). Thus, END repopulation occurs due to the unprecedented level of microglia depletion rather than drug treatment. Together, these data provide evidence for the existence of a very small population of myeloid cells located in the SVZ and adjacent WM tracts that can uniquely survive sustained high dose CSF1Ri treatment.

### Empty microglial niche-dependent (END) repopulation elicits a dynamic wave of repopulating proliferative myeloid cells

In PND repopulation, microglia repopulate the brain parenchyma in a homogeneous fashion, with repopulating cells displaying no preference for specific locations (Figure 1C). In contrast, END repopulating cells first appear in precise ventricular and WM locations (Figure 1D2), similar to the locations of the few surviving myeloid cells. To expand upon this initial observation, we sought to define the anatomical niches of this distinct form of repopulation and built a spatial heat map at three brain positions (Bregma 2.58 mm, 1.10 mm, -2.06 mm) along the rostral-caudal axis in brains of mice at 7d recovery (Figure 3A). Throughout this axis, IBA1^+^ cells initially repopulate the brain within the caudoputamen, particularly in areas near the lateral ventricle and associated WM tracts. At rostral regions of the brain, cells are found within the rostral migratory stream (RMS), a projecting axonal tract from the SVZ to the olfactory bulb. In more caudal brain regions, repopulating cells are seen near the SVZ, caudoputamen, and corpus callosum. Subsequent analysis of the entire brain confirms the presence of these early repopulating cells in areas near the SVZ/ventricular zones, WM tracts, and caudoputamen.

**Figure 3.**
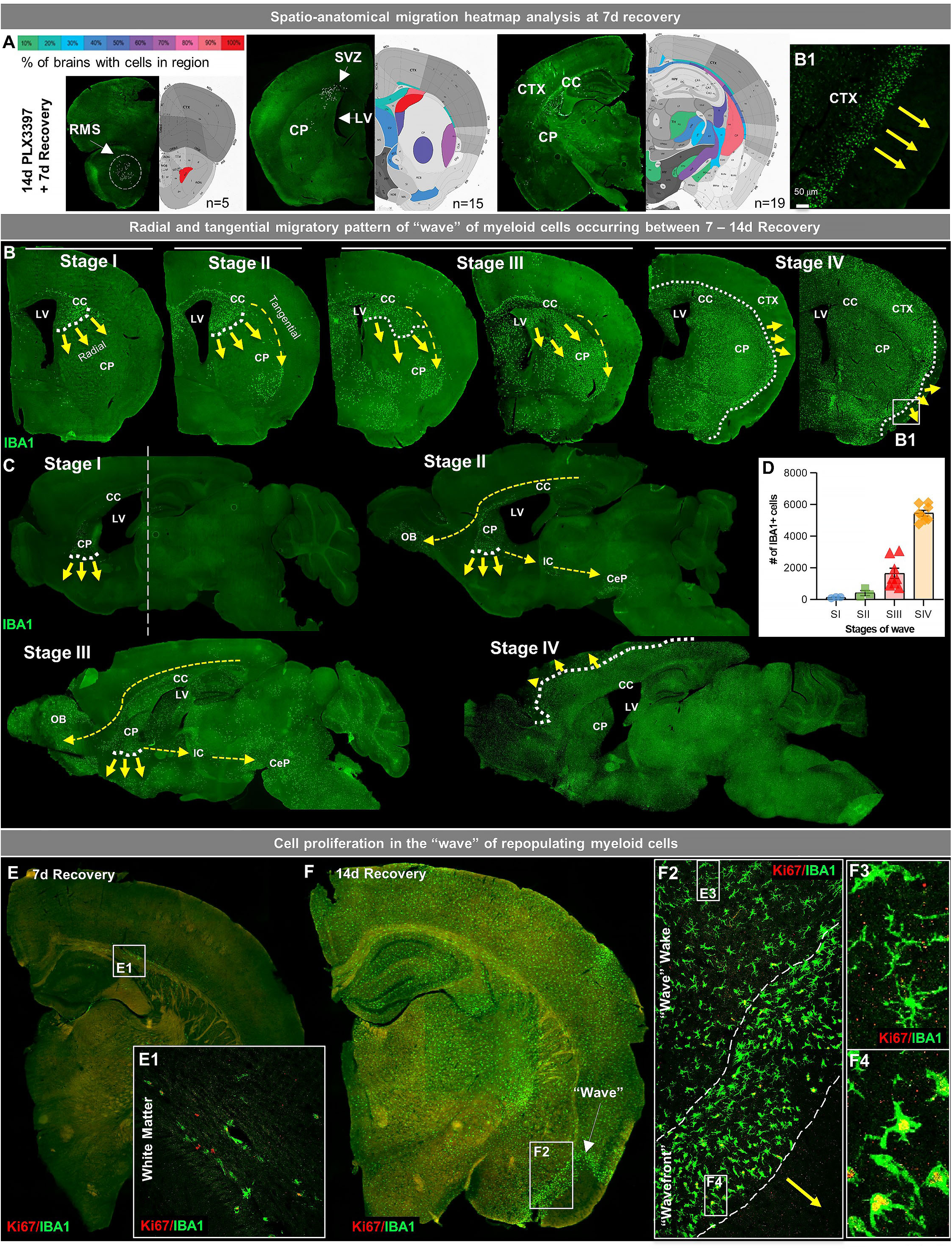
**Empty microglial niche-dependent (END) repopulation elicits a dynamic wave of repopulating proliferative myeloid cells.** (A) Spatial heat map of END repopulation at 7d recovery in three brain positions (Bregma 2.58 mm, 1.10 mm, -2.06 mm) depicting % of brains with cells present in specific brain regions (n=5-19). (B-C) Immunofluorescence whole brain coronal (B) and sagittal (C) section images of IBA1^+^ cells (green) show a sequential time course of the “wave” (see higher resolution image of wavefront in B1 insert) of myeloid cells filling the brain between 7d and 14d recovery during END repopulation. White dotted lines and yellow arrows indicate the edge and direction of the “wave”, highlighting the radial migratory patterns of END repopulating cells. Yellow dashed arrows indicate the direction of the tangential migratory pattern of END repopulating cells, mostly utilizing WM tracts. The straight white dashed line in C shows the Bregma position at which coronal sections were taken for B. (E-F) Representative immunofluorescence whole brain images of myeloid cells (IBA1, green) and proliferating cells (Ki67, red) at 7d (A) and 14d (B) recovery. Insets show higher resolution of IBA1 and Ki67 colocalization in initially repopulating cells (A1) and in cells outside and inside the wavefront (B2-4). Data are represented as mean ± SEM (n=3-8).

Repopulating cells migrate in a radial and tangential fashion, initially filling the WM and caudoputamen before spreading out through the cortex in a dynamic wave between 7d and 14d recovery (Figure 3B-C). Analysis of the migratory wave during this recovery time from Bregma 0.445 to Bregma -0.08 shows repopulating cell deposition appears to occur in stages. At Stage I, cells are visible near the lateral ventricle at the intersection of the corpus callosum, caudoputmen and SVZ, and appear to migrate in a radial migration pattern from the ventricular zone into and filling the caudoputamen in an inferior direction. At this stage, cells also appear in WM tracts, specifically in the corpus callosum. At Stage II, cells begin to migrate in a tangential migration pattern utilizing WM tracts to migrate into WM areas or areas near WM tracts. At Stage III, repopulating cells continue to migrate in both a radial and tangential migratory pattern filling ∼80-95% of the striatum, including the caudoputamen, lateral septal complex, and pallidum. It is also apparent that repopulating cells are restricted or unable to migrate past the WM tract or corpus callosum between the cerebral nuclei and cerebral cortex. At Stage IV, the cells have broken through this WM tract barrier and migrate out in a radial migration pattern moving from the subcortical to the cortical plate (Figure 3B-D).

In contrast to the proliferative profile of previously described PND-repopulating microglia, in which remaining microglia proliferate throughout the brain to give rise to newly repopulating cells, proliferating END repopulating myeloid cells are initially only found near ventricles and WM tracts (Figure 3E, 3E1). As the cells spread and migrate through the brain, proliferation continues but remains localized within the cell “wavefront” (Figure 3F, 3F2, 3F4). Once out of the front (i.e. in the wake of the wave) IBA1^+^ cells appear to stop proliferating (i.e. Ki67^-^; Figure 3F3). This wave of proliferating cells is most apparent at Stage IV, led by a wavefront with an average width of ∼100-150 µm of amoeboid and proliferating myeloid cells.

### END repopulating myeloid cells originate from an existing *Cx3cr1+* cell source originating from the SVZ/WM area

Since repopulating cells first appear in the SVZ and the SVZ is a notable neurogenic/proliferative niche in the brain, we next stained sections containing the SVZ in control, 14d PLX3397, 5d and 7d recovery groups for known precursor cell markers, as well as other cell lineage markers. Between 3 - 5d recovery, repopulating cells transiently express NESTIN (Figure 4A), MASH1 (Figure 4B), and TIE2 (Figure S2A), but are negative for GFAP (Figure S2B), DCX (Figure S2C), OLIG2 (Figure S2D), and SOX2 (Figure S2E) at all timepoints. Consequently, we performed lineage tracing using tamoxifen-inducible *Cre*-recombinases under control of the *Nestin* (*Nestin^CreERT2^*) and *Ascl* (*Ascl1^CreERT2^*, note: *Ascl1* encodes for MASH1) promoters, along with the myeloid cell-specific line (*Cx3cr1^CreERT2^*). *Cre* lines were crossed with YFP reporters for visualization of induced expression (Figure S2F). Tamoxifen was given immediately following PLX3397 treatment to track the lineage of repopulation cells, except for *Cx3cr1^CreERT2^* mice - which were given a 21d washout period (i.e. tamoxifen was administered 21d prior to PLX3397 treatment) to restrict labeling to resident vs short-lived peripheral myeloid cells^36^. These studies revealed that repopulating cells do not originate from *Ascl1*^+^ (Figure 4C) or *Nestin*^+^ (Figure 4D) cell sources, despite their transient expression of these markers. Consistent with previous reports^37, 38^, we found that the *Cre*-recombinase from the *Cx3cr1^CreERT2^* line is leaky in the absence of tamoxifen (∼25% of microglia express YFP; Figure S2G-H). Despite this, we show that ∼100% of microglia, and repopulating cells, expressed YFP following tamoxifen administration 3 weeks prior to microglial elimination, thereby demonstrating that repopulating cells derive from a *Cx3cr1*^+^ cell source (Figure 4E). These repopulating cells exhibit a similar wave-like migration pattern, appearing first near ventricular areas and lastly in cortical regions (Figure S2I).

**Figure 4.**
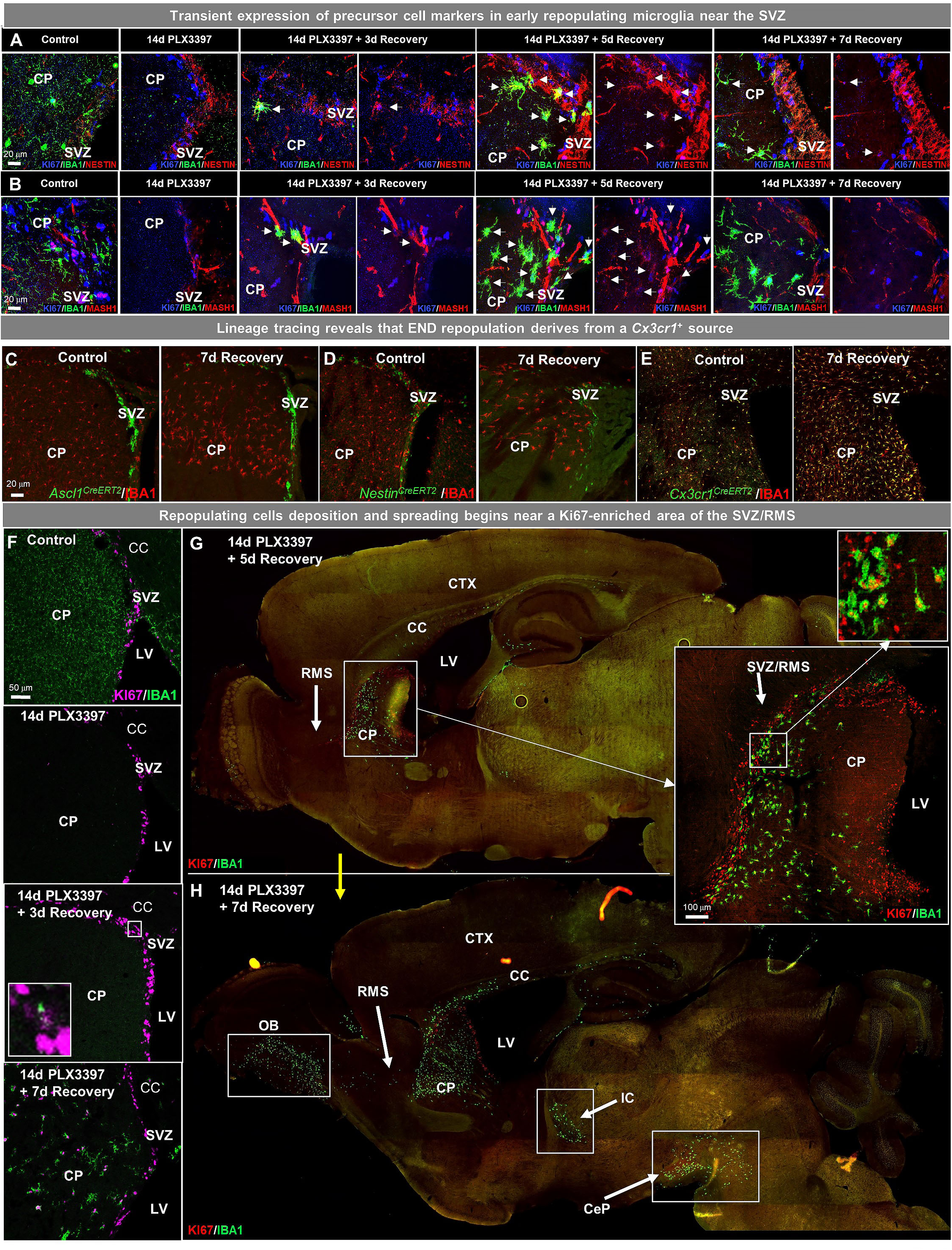
**END repopulating myeloid cells originate from an existing *Cx3cr1+* cell source originating from the SVZ/WM area:** (A-H) Two-month-old WT mice were treated with PLX3397 (600 ppm) for 14d, then allowed to recover without PLX3397 for 3, 5, 7 days. (A-B) Representative 63x immunofluorescence images of proliferating (Ki67^+^, blue) myeloid cells (IBA1^+^, green) staining for positive for common cell lineage/precursor cell markers: NESTIN (red, A) and MASH1 (red, B) in the SVZ of control 14d PLX3397, 3d recovery, 5d recovery, and 7d recovery mice. (C-E) CreER-directed lineage-specific labeling. In these mouse lines, tamoxifen-inducible Cre-recombinase is expressed under control of the promoter of interest. When activated by tamoxifen, the CreER fusion protein translocates to the nucleus allowing transient recombination to occur and, when crossed to a YFP reporter, visualization of induced expression via eYFP. (C-D) Representative 20x images of IBA1^+^ (red) and associated promoter-driven lineage-derived (YFP, green) cells in control and 7d recovery *Ascl1^CreERT2^/YFP* (B) and *Nestin^CreERT2^/YFP* (C) mice. (E) Representative 20x images of IBA1^+^ (red) and *Cx3cr1^+^* lineage-derived (YFP, green) cells in control and 7d recovery mice. (F-H) Representative coronal (F) and sagittal (G-H) brain images of IBA1^+^ (green) and Ki67^+^ (red, B-C) cells near SVZ/RMS regions between 5-14d recovery. Inserts provide higher resolution images of cells near the SVZ/RMS proliferative site of repopulation, illustrating the spread of repopulating cells via WM/axonal tracts (i.e. RMS) between the CP and OB and between other WM regions (IC and CeP). CP, caudoputamen; CC, corpus callosum; CTX, cortex; LV, lateral ventricle; RMS, rostral migratory stream; SVZ, subventricular zone; OB, olfactory bulb; IC, internal capsule; CeP, cerebral peduncle.

### The role of the SVZ/WM area in myeloid cell proliferation and migration signaling during early END repopulation

In addition to the site of surviving CSF1Ri-resistant microglia, we next explored what role the SVZ/WM area plays in cell proliferation and the spatio-temporal expansion of repopulating cells during END repopulation. Immunohistochemical analyses for IBA1 and Ki67 show that repopulating cells initially appear between 3d - 7d recovery, with IBA1^+^/Ki67^+^ cells apparent within the SVZ and the adjacent caudoputamen by 7d recovery (Figure 4F). Further evaluation of sagittal sections from mice at 5d recovery confirms that repopulating cells first populate the parenchyma in the SVZ, specifically from a Ki67^+^-dense region located along the alpha arm of the SVZ (αSVZ) and posterior RMS (pRMS) (Figure 4G). These cells subsequently accumulate in the WM areas adjacent to the SVZ (i.e. the corpus callosum), caudoputamen, RMS, olfactory bulb, internal capsule, and cerebral peduncle before eventually filling the parenchymal grey matter (Figure 4H). The internal capsule is a WM structure/projecting axonal tract that connects the CP to the cerebral peduncle, while the RMS connects the SVZ to the olfactory bulb, providing anatomical pathways by which repopulating cells travel to specific brain locations. Of note, the aforementioned niches are the precise locations in which microglia initially colonize the developing brain^7, 10, 12^. During the 14^th^ – 17^th^ week of gestation in the developing human brain, microglia are found near or within: the optic tract, the WM junction between the thalamus and internal capsule, and the junction between the internal capsule and the cerebral peduncle^39^.

To examine the transcriptional changes occurring in the SVZ during this early stage of END repopulation, we micro-dissected the SVZ from control, 14d PLX3397, and 5d recovery mice, and performed bulk tissue gene expression analyses via RNA-seq (Figure 5A). Gene expression data can be explored at http://rnaseq.mind.uci.edu/green/alt_repop_svz/gene_search.php. In comparing control vs. 5d recovery, 227 DEGs were identified (FDR < 0.05) between the two groups (Figure 5B), with the majority being downregulated microglia-enriched/related genes, reflecting the reduced pool of myeloid cells in the CNS during the early stages of repopulation. Upregulated non-myeloid enriched DEGs in depleted vs. 5d recovery mice (Figure 5B) consisted of genes implicated in cell cycle regulation (*Pak3, Swi5, Psmd11, Stat3*), DNA transcription/recombination/repair/expression (*Alyref2, Swi5, Zfp612, Zfp51, Thumpd1, Prmt5, Taok3, Psmd11, Tox, Stat3*), cell adhesion/migration/proliferation (*Pak3, Anxa1, Cadm1*) and development (*Gfap, Rab14, Zfp612*). Gene ontology (GO) analysis of DEGs between control and 5d recovery SVZ tissue identified the following top four enriched pathways: *myeloid cell differentiation, leukocyte immunity, leukocyte activation, leukocyte chemotaxis and phagocytosis* (Figure 5C). Focusing on myeloid genes, *P2ry12*, *Siglech*, *Trem2*, *Cd33* and *Cx3cr1* were least enriched during initial repopulation, whereas *Ccl12*, *Cd52*, *Lyz2*, *Itgb2*, and *Cd84* were highly enriched (Figure 5D). To explore the biological relevance of these findings and the effect on early repopulation dynamics due to a loss in one of these important genes/signals, we administered an antibody against CCL12, the most highly upregulated gene during early END repopulation (Figure 5E). Here, we show that neutralization of CCL12 results in a significant reduction in repopulating cell numbers at 7d recovery, but not total distance of cell spreading (Figure 5F-H), indicating that this chemokine may play an important role in early repopulating cell proliferation. Together, these data highlight the role of the SVZ and signaling during early END repopulation.

**Figure 5.**
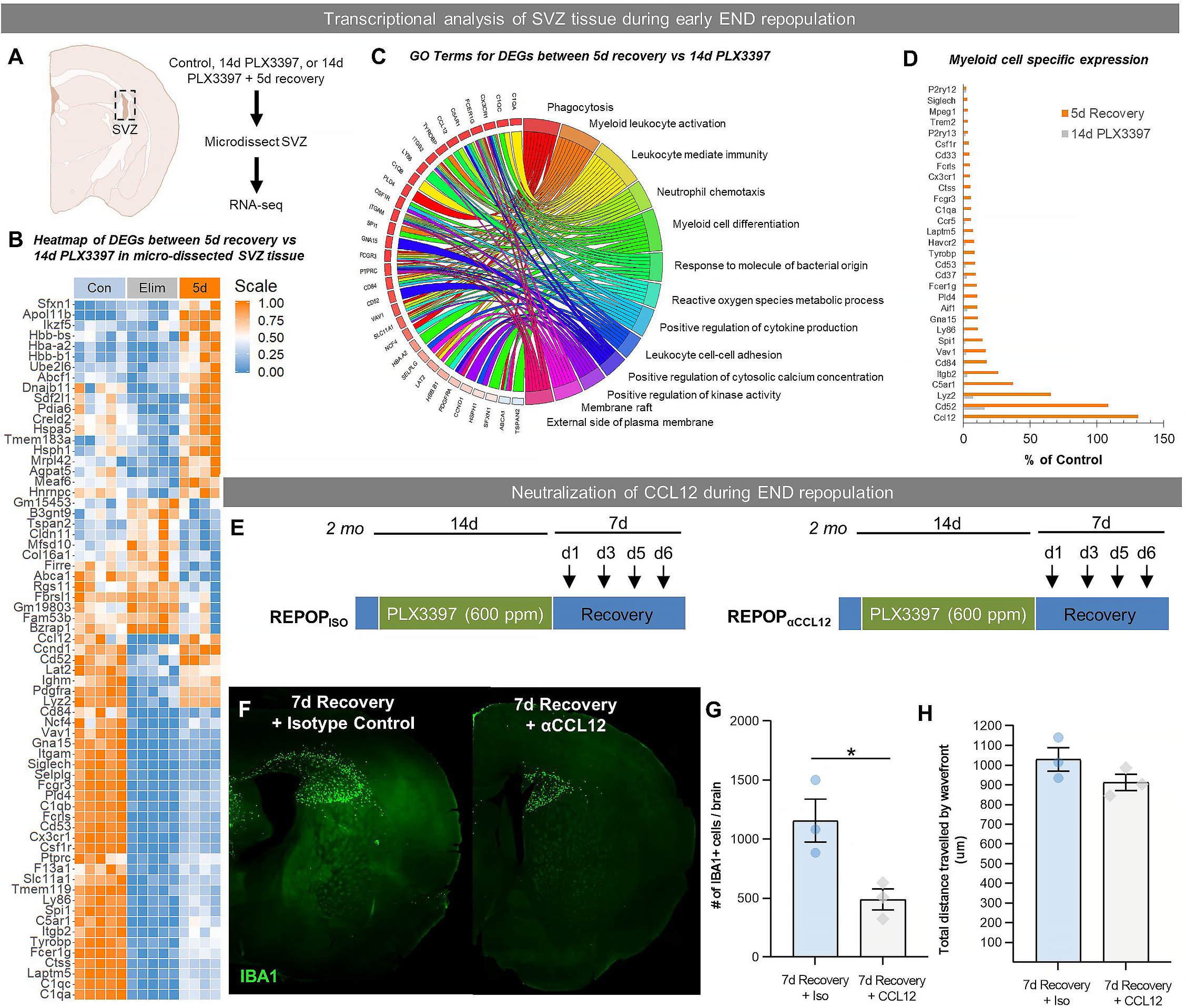
**The role of the SVZ/WM area in myeloid cell proliferation and migration signaling during early END repopulation.** (A-D) Transcriptional analysis of SVZ tissue during early END repopulation. (A) Bulk tissue RNA-seq analysis was performed on micro-dissected SVZ tissue from control, 14d PLX3397, and 5d recovery brains (n=5). (B) Heatmap of DEGs between 14d PLX3397 (Elim) and 5d recovery SVZ tissue. Gene expression data can be explored at http://rnaseq.mind.uci.edu/green/alt_repop_svz/gene_search.php. (C) Gene ontology chord plot of DEGs between control and 5d SVZ tissue. (D) Plot highlighting expression (% of control) changes in myeloid-associated genes in depleted (14d PLX3397) and repopulated (5d recovery) SVZ tissue. (E-H) Neutralization of CCL12 during END repopulation. (E) Experiment schematic of CCL12 neutralization study: 2-month-old WT mice were treated for 14d with PLX3397 (600 ppm) and then placed on control diet for 7d allowing for END repopulation. Four i.p. injections were administered of anti-CCL12 antibody or goat IgG (Isotype control) at 1d recovery, 3d recovery, 5d recovery and 6d recovery. (F) Representative whole brain images of IBA1^+^ cell (green) deposition following treatment. (G-H) Quantification of number of total IBA1^+^ cells and total distance traveled by IBA1^+^ wavefront in (F).

### END repopulating cells do not derive from the periphery

Our data show that extensive microglial depletion results in repopulation of the adult brain from myeloid cells that originate from SVZ/WM areas and utilize WM “highways” to spread throughout the brain before filling the cortex in a distinctive wave-like pattern. As END repopulating cells maintain distinct phenotypes from microglia, even after extended periods of time in the brain, we concluded that either 1) surviving SVZ/WM myeloid cells represent a distinct myeloid cell type with the capacity to spread and fill the empty microglial brain niche, or 2) extensive microglial depletion stimulates an influx of peripheral cells, which enter near the ventricles and then spread throughout the brain maintaining distinct profiles to their microglial counterparts. Indeed, previous studies have shown that under certain conditions (e.g. in an empty microglial niche) induced by microglial depletion, peripheral myeloid cells can infiltrate and serve as the source for microglial repopulation in the brain parenchyma^40–43^. Thus, we reasoned that repopulation could be occurring from peripheral sources and undertook several complementary experimental approaches to explore this.

The choroid plexus is a major route of cellular entry into the CNS^44^, thus we explored the contribution of this site to END repopulation. Here, we observe that choroid plexus myeloid cells do not repopulate until 14d recovery, despite the appearance of myeloid cells in the adjacent brain parenchyma (Figure 6A-B). Utilizing *Cx3cr1*^GFP/+^/*Ccr2*^RFP/+^ mice, in which CCR2^+^ cells (mainly monocytes) express RFP, we show that END repopulating cells are CCR2/RFP^-^, or not a result of the infiltration of CCR2^+^ monocytes (Figure S3A-C). A recent study has posited that CSF1R inhibition suppresses CCR2^+^ monocyte progenitor cells and CX3CR1^+^ BM-derived macrophages (among other BM populations) and that these populations do not recover after cessation of CSF1R inhibition^45^. In this study, we evaluated the effects of 14d PLX3397 600 ppm on peripheral myeloid cells, including CCR2^+^ and CX3CR1^+^ myeloid cells, and BM myeloid precursors (Figure S3D-F). Here, we observe an expansion, not suppression, of HSCs, common myeloid progenitors (CMPs), and granulocyte/monocyte progenitors (GMPs) following CSF1R inhibition and/or recovery (Figure S3F). However, these changes do not result in significant changes to blood or spleen myeloid cell populations (Figure S4D-E). CCR2/CCL2 signaling is implicated in many neuropathologies with peripheral cell CNS infiltration^46^, however, we show that CX3CR1 and CCR2 KO in *Cx3cr1*^GFP/GFP^/*Ccr2*^RFP/RFP^ mice elicits no alterations in END repopulation (Figure 6C-D) indicating that this form of repopulation is not dependent on these signaling axes. We further explored the CCR2/CCL2 signaling axis using *Ccl2*^-/-^ mice. Interestingly, we found that a lack of *Ccl2* conveyed resistance to CSF1Ri-induced microglial death, with 14d PLX3397 treatment only eliminating ∼90-95% of microglia (Figure S3G-H). As a result, PND repopulation was observed in these mice rather than END repopulation. In combination with findings from H2K-BCL2 mice, these data confirm that >99% microglial elimination is a requirement for END repopulation, which may not be achieved by 14d PLX3397 treatment under certain conditions.

**Figure 6.**
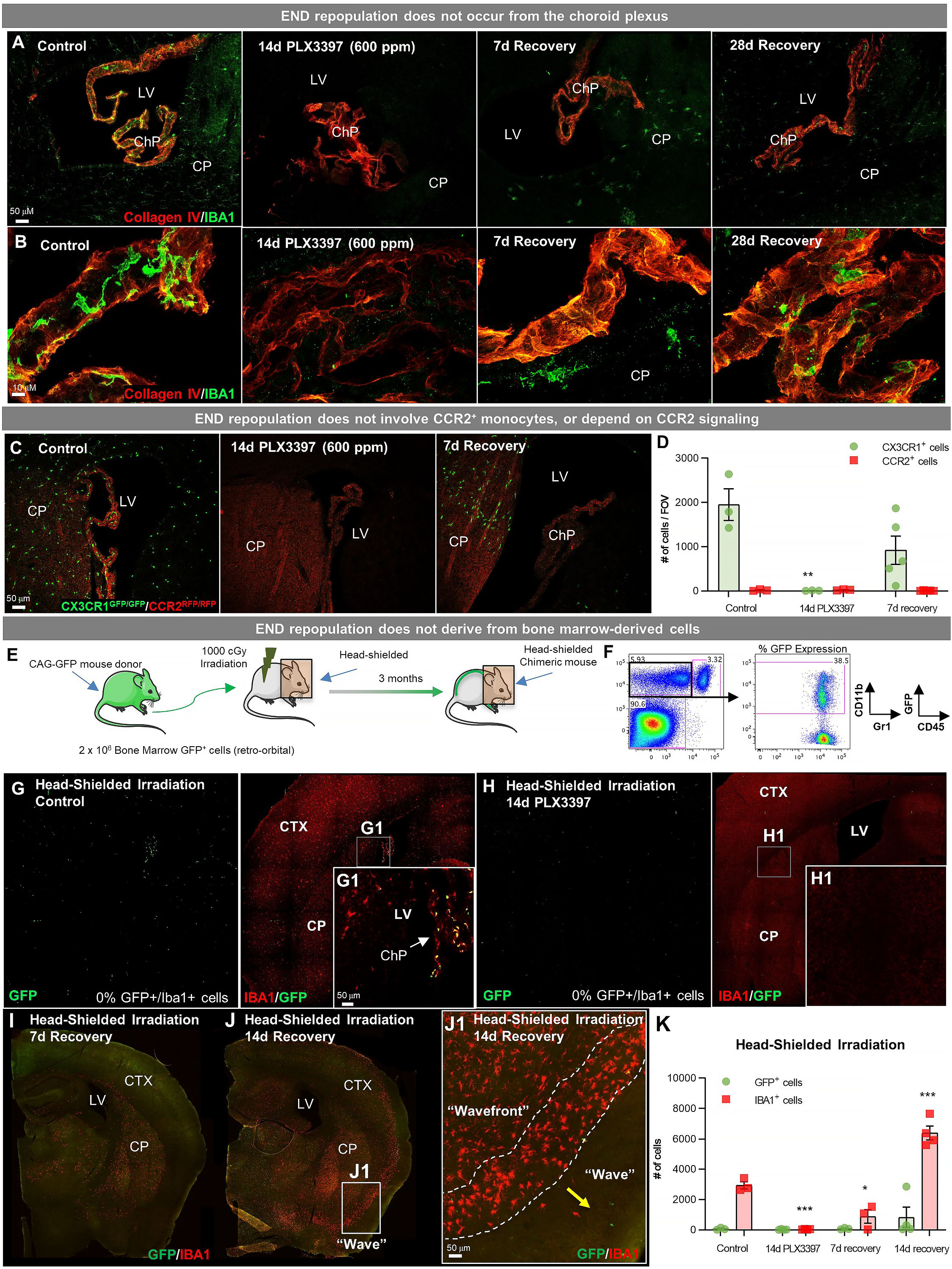
**END repopulating cells do not derive from the periphery.** (A-B) Representative immunofluorescence 10x (A) and 63x (B) images of IBA1^+^ cell deposition within the choroid plexus (labeled with Collagen IV) in control, 14d PLX3397, 7d recovery, and 28d recovery mice. (C, D) Sustained microglial depletion and END repopulation in *Cx3cr1*^GFP/GFP^/*Ccr2*^RFP/RFP^ (i.e. *Cx3cr1* and *Ccr2* KO). Representative immunofluorescence images (C) and quantification of the number (D) of CX3CR1-GFP^+^ (green) and CCR2-RFP^+^ (red) cells in control 14d PLX3397, and 7d recovery mice (n=3-5). (E) Experimental paradigm: Schematic depicting generation of BM GFP^+^ chimeras, achieved by head-shielded (HS) irradiation and transplantation of donor GFP+ BM cells. After 3 months mice were treated for 14d with PLX3397 and then allowed to recover for 7d or 14d on control diet. (F) FACS gating strategy to determine % chimerism in BM GFP^+^ chimeras achieved by head-shielded irradiation. (G-J) Representative whole brain images of GFP^+^ (green) and IBA1^+^ (red) cell deposition in HS irradiated control (G), 14d PLX3397 (H), and mice following 7d (I) and 14d recovery (J). (J1) Higher resolution images of repopulating cell wavefront seen in HS chimeras during END repopulation. (K) Quantification of number of GFP+ and IBA1+ cells treated HS irradiated mice. Data are represented as mean ± SEM (n=3-7). *p < 0.05, ** p < 0.01, *** p < 0.001. CTX, cortex; CP, caudate putamen; LV, lateral ventricle; ChP, choroid plexus.

Next, we utilized bone marrow (BM) chimeric mice to further determine whether repopulating cells originate from a peripheral source (i.e. non-CCR2+ monocytes or other BM-derived cells). Two-month-old wildtype mice underwent head-shielded (HS) irradiation, followed by retro-orbital administration of GFP^+^ donor BM cells and 12 weeks of recovery for immune system reconstitution (Figure 6E-F). Previous studies have shown that CNS infiltration can occur from the BM upon exposure to head irradiation and consequent BBB permeability^47–49^. With HS irradiation, however, no GFP^+^ cells were visible in the parenchyma of control chimeric mice (Figure 6G, K), confirming that under normal conditions circulating peripheral cells do not enter the brain. GFP^+^ cells were visible in the choroid plexus of control HS chimera (Figure 6G1), consistent with their partial turnover by circulating BM-derived cells^50^. Fourteen-day PLX3397 eliminated all myeloid cells in HS-irradiated brains (Figure 6H, K). By 7d recovery, END repopulation was apparent in HS-irradiated chimeric mice, however, all IBA1^+^ cells were GFP^-^ (Figure 6I-K), thus ruling out peripheral BM-derived cells as the source of this form of microglial repopulation. At 14d recovery, the wave of repopulating myeloid cells was visible in the cortex, as seen in WT mice (Figures 6J, 6J1). It should be noted that since all parenchymal repopulating IBA1^+^ cells were GFP^-^ in HS chimeric mice, these data provide strong evidence that BM-derived cells, including choroid plexus macrophages - which are GFP^+^ in HS chimeric control mice - do not contribute to END repopulation. Together, these data indicate that surviving SVZ/WM myeloid cells serve as the major source of this unique form of CNS myeloid cell repopulation.

### Repopulating myeloid cells are transcriptionally distinct and mount a differential response to inflammatory stimulus compared to homeostatic microglia

We next sought to gain insight into the transcriptional profile of END repopulating cells once in residence in the brain. In addition, we also investigated whether phenotypic and transcriptional alterations translated into functional consequences and thus explored their response to immune challenge, via LPS administration (Figure 7A). Here, we performed RNA sequencing (RNAseq) on FACS-sorted CD11b^+^CD45^int^ control and 28d recovery cells at 6 and 24 hr following LPS-induced immune challenge (Figure 7A, S4A). For controls, cells were also collected from mice that did not receive LPS, referred to as 0 hr post LPS. Flow cytometry analysis of CD11b^+^CD45^int^ cells collected at 28d recovery shows that these repopulating cells exhibit higher CD45 and CD11b intensities compared to control microglia at baseline. Control microglia show increased CD45 and CD11b in response to LPS, while repopulated cells do not change (Figure S4B-C).

**Figure 7.**
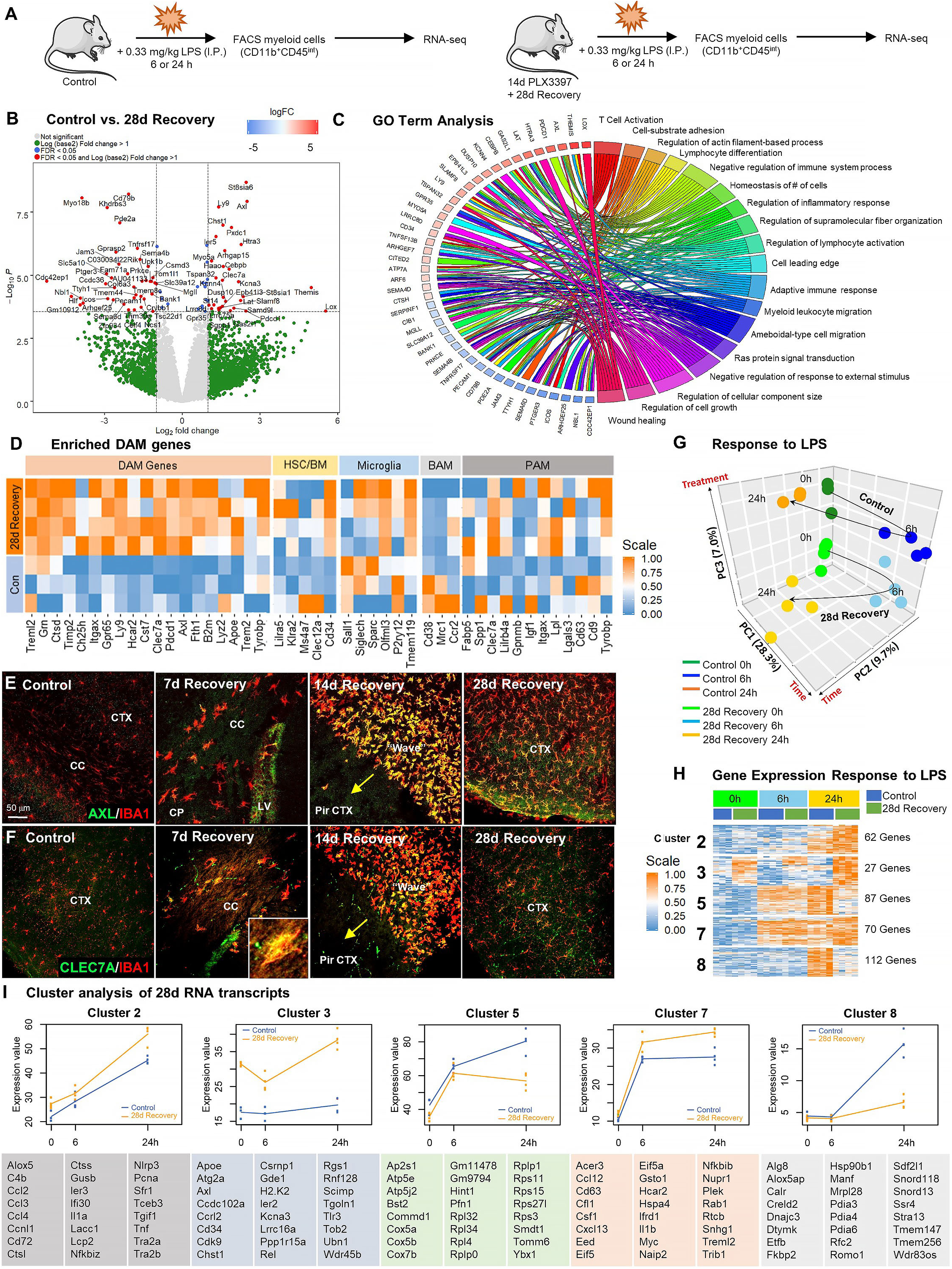
**Repopulating myeloid cells are transcriptionally distinct and mount a differential response to inflammatory stimulus compared to homeostatic microglia.** (A) Two-month-old WT control or 28d recovery mice were given intraperitoneal injections of either PBS or LPS (0.33mg/kg) and then collected at 6 or 24 hours post injection. Controls, which were mice that did not receive LPS, are referred to as 0 hr post LPS. Myeloid cells were extracted from whole brain hemispheres, isolated using FACS gating for CD11b^+^CD45^int^ and processed for RNA-seq. (B) Volcano plots displaying the fold change of genes (log2 scale) and their significance (y axis, -log10 scale) between control vs. 28d recovery mice. (C) Gene ontology chord plot of DEGs between control and 28d recovery myeloid cells. (D) Heatmap showing expression of genes enriched in DAM, HSC/BM-derived cells, canonical microglia, BAM, and PAM signatures in control and 28d repopulating myeloid cells. (E-F) Representative immunofluorescence 20x images of IBA1^+^ (red) and AXL^+^ (green, I) or CLEC7A^+^ (green, J) cells shown in areas with high repopulating cell deposition in control, 7d, 14d, and 28d recovery mice. (G)Principal component analysis plot of extracted control and 28-day recovery cells, across time (0h, 6h, 24h) and treatment (+/- LPS), depicting the separation of groups into six clusters. (H) Heatmap of selected time-series cluster analysis of control and 28d recovery cells. Provided number indicates number of genes per cluster. (I) Time-series cluster analysis of control vs. 28d recovery myeloid cell response (during END repopulation) to LPS challenge following 14d PLX3397 (600 ppm in chow; from H). Clusters showing distinct responses to LPS between control and 28d END repopulated cells, across time, were plotted as eigengene values, along with the top represented genes within each cluster. Data are represented as mean ± SEM (n=3-5). *p < 0.05, ** p < 0.01, *** p < 0.001. CP, caudoputamen; CC, corpus callosum; CTX, cortex; LV, lateral ventricle; PirCTX, piriform cortex. Gene expression data can be explored at http://rnaseq.mind.uci.edu/green/alt_repop_lps/gene_search.php.

RNA was extracted from FACS-sorted CD11b^+^CD45^int^ cells and RNA-seq analysis was performed to establish a high-resolution transcriptome profile of these cells in the absence and presence of LPS. Gene expression data can be explored at http://rnaseq.mind.uci.edu/green/alt_repop_lps/gene_search.php. Unlike PND repopulation, in which control and repopulating cells display few transcriptional differences (Fig. S4F-G), we identified 69 DEGs in END repopulated myeloid cells compared to control microglia in the absence of LPS (logFC > 1, FDR > 0.05; Figure 7B). GO analysis of DEGs between control and 28d repopulated cells identified the following top five enriched pathways: *regulation of cellular component size, negative regulator of chemotaxis, ameboidal-type cell migration, negative regulation of response to external stimulus*, and *wound healing* (Figure 7C). To visualize differences in these cells vs. other myeloid cell subsets, we compared their gene expression profile to previously established myeloid cell signatures, including DAMs/MGnD/ARMs^15–17, 51^, PAMs^20^, border-associated myeloid cells (BAMs)^52^/CNS-associated macrophages (CAMs)^53^, HSC/BM-derived myeloid cells^54^, and homeostatic microglia^55, 56^. Notably, we observe robust enrichment of DAM-associated genes in repopulating cells, including *Clec7a*, *Axl*, *ApoE*, *Cst7*, *Ctsd*, and *Ly9* (Figure 7D). AXL and CLEC7A immunostaining is apparent in repopulating myeloid cells, particularly in the early stages of repopulation, and in cells located at the wavefront, whereas undetected in microglia from control brains (Figure 7E-F). In addition, we also observe a reduction in expression of *Sall1*, a transcription factor unique to microglia^57^, in repopulating compared to homeostatic microglia.

After confirming distinct transcriptome differences between control and 28d recovery isolated microglia at 0h LPS (i.e. in the absence of LPS), we next evaluated their gene expression profiles at 6h and 24h following LPS challenge. Principal component analysis demonstrated that biological replicates were highly correlated, with samples clustering into six distinct groups (Figure 7G). To identify global patterns in gene expression changes over time between our experimental groups, we employed K-means clustering^58^, revealing 9 distinct clusters of genes (Figure 7H, S4H). Cluster 3 contains genes that are significantly different between control and repopulating cells across all time points, including genes implicated in clearance (*Apoe*, *Atg2a*, *Axl, Wdr45b)*, cell growth/differentiation (*Cd34, Cdk9)*, stress (*Ier2, Ppp1r15a)*, inflammation (*Ccrl2, H2.K2, Scimp, Tlr3)* and senescence (*Ubn1, Wdr45b).* Clusters 2 (e.g. *C4b, Ccl2, Ccl3, Ccl4, Ctsl, Ctss, Il1a, Nlrp3, Pcna* and *Tnf*), 5, 7 (e.g. *Ccl12, Cd63, Csf1, Cxcl13, Il1b, Myc, Plek* and *Nfkbib*), and Cluster 8 shows differential responses to LPS at the 24h timepoint between control and repopulating cells (Figure 7I). GO term analysis of Cluster 3 revealed top enriched pathways: *phagopore assembly site membrane*, *extrinsic component of membrane*, and *endosome*. Cluster 8 GO term analysis revealed the following enriched pathways: *negative regulation of glucocorticoid secretion*, *negative regulation of interleukin-1 mediated signaling pathway*, and *connective tissue replacement involved in inflammatory response wound healing*. Together, these findings demonstrate that END repopulating cells represent a myeloid cell population that are transcriptionally and functionally distinct from adult homeostatic microglia.

### Repopulating myeloid cells elicit few functional differences in behavior, injury, and neuronal-associated structures

Given the transcriptional and functional distinction between END and homeostatic microglia, we next sought to determine the physiological and functional consequences of filling the adult brain with these cells. Previous studies have shown that replacement of endogenous microglia with PND repopulating microglia results in no detectable changes in cognitive or behavioral function^32, 59^. Here, we performed a battery of cognitive and behavior tasks in 28d recovery mice (Figure 8A-H), and observe that these mice exhibit reduced locomotion (Figure 8A-B) and reduced ability to discriminate a novel place change compared to controls (Figure 8H). However, we observe no other significant behavior or cognitive disruptions. Overall, these animals do not appear to exhibit overt cognitive, behavioral, or health changes.

**Figure 8.**
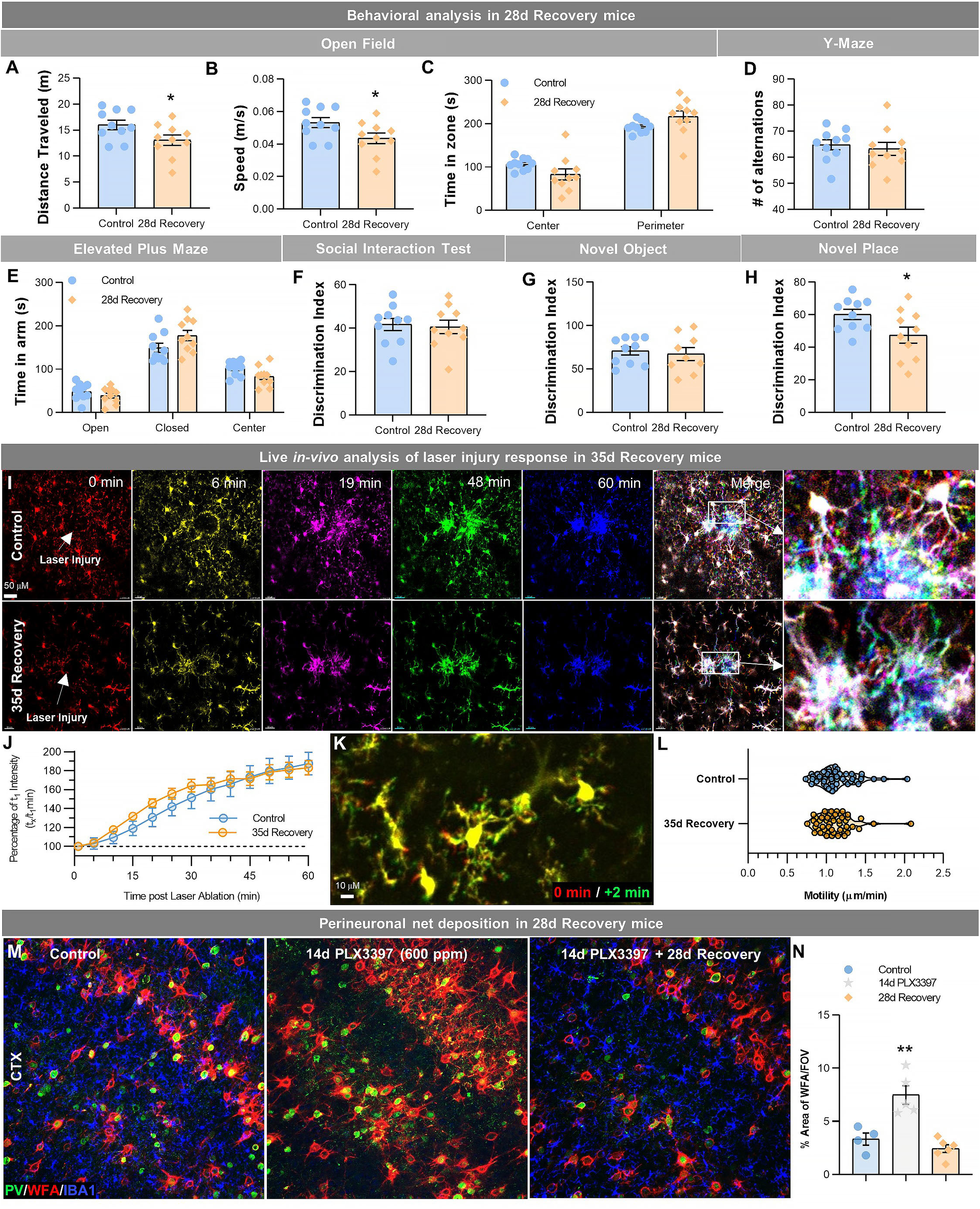
**Repopulating myeloid cells elicit few functional differences in behavior, injury, and neuronal-associated structures.** (A-H) Two-month-old WT mice treated with vehicle or PLX3397 (600 ppm in chow) for 14 days, followed by 28 days of recovery, filling the brain with alternatively repopulated myeloid cells. Mice underwent behavioral assessment by Open Field, Y-Maze, and Elevated Plus Maze, Social Interaction Test, and Novel Object/Place recognition. In Open Field, distance traveled (A) and average speed (B) were reduced in 28d recovery groups, but time spent in each zone (C) was unchanged. No changes in performance were seen in the Y-maze, as measured by the number of alternations (D). No changes in anxiety behaviors were seen in the elevated plus maze, as measured by time spent in the open or closed arms (E). No changes in social preference were seen in the social interaction test (F). No changes in novel object recognition memory were seen (G), but 28d recovery mice had a significant impairment in novel place recognition memory (H). Data are represented as mean ± SEM (n=10). *p < 0.05. (I-L) 2-month-old Cx3cr1-GFP^+^ mice treated with vehicle or PLX3397 (600 ppm in chow) for 14 days, followed by 35 days of recovery, filling the brain with alternatively repopulated myeloid cells. (I) Analysis of motility and focal laser response in alternatively repopulating myeloid cells. Representative images of Cx3cr1-GFP^+^ myeloid cell response to laser ablation, over time, obtained via two-photon imaging in control and 35d recovery mice. (J) Quantification of the average normalized GFP^+^ intensity measured within a 50 µm radius of the site of damage over time. (K) Representative image of Cx3cr1-GFP^+^ myeloid cell at process motility from 0 min (red) to 2 min (green) time-period. (L) Quantification of process motility (i.e. extension of process µm per min) measuring the difference in visibly moving processes over 2 min in control and 28d recovery myeloid cells. Data are represented as mean ± SEM (n=4). (M-N) 2-month-old WT mice treated with vehicle or PLX3397 (600 ppm in chow) for 14 days, followed by 28 days of recovery, filling the brain with alternatively repopulated myeloid cells. (M) Representative immunofluorescence 20x images of Parvalbumin^+^ (PV, green), WFA (a marker for perineuronal nets, red), and IBA1^+^ (blue) cells shown in the cortex of control, 14d PLX3397, and 28d recovery mice. (N) Quantification of % area of WFA per field of view (FOV) in the cortex. Data are represented as mean ± SEM (n=4-6). ** p < 0.01.

Based on transcriptional changes in myeloid signaling and priming, we next explored how these repopulating cells would respond to injury. Live two-photon imaging of cortical myeloid cells was performed in control and 35d recovery Cx3cr1^GFP/+^/Ccr2^RFP/+^ mice (Figure 8I-L). Cx3cr1-GFP^+^ repopulating myeloid cells react to a focal laser injury with similar rates of migration as homeostatic microglia (Figure 8J, Supplemental videos S1 and S2). Furthermore, extension and retraction of processes (i.e. motility) were similar between control microglia and repopulating cells, although repopulating cells displayed fewer and thicker processes as previously demonstrated (Figure 8K-L).

Recent studies have identified a novel contribution of microglia in modulating perineuronal nets (PNNs) in the adult brain^60, 61^, specialized extracellular matrix assemblies that enwrap neurons and proximal dendrites to regulate synaptic plasticity^62^, protect against neurotoxins^63^, and enhance signal propagation^60, 61^, among other functions. In control brains, PNNs (as detected by *Wisteria floribunda* agglutinin (WFA) staining) are preferentially found on parvalbumin (PV) expressing cells. Fourteen-day treatment of high dose PLX3397 results in significant elevation in PNN staining, corroborating previous findings that microglia play a critical homeostatic role in modulating these structures^60^. Following CSF1Ri withdrawal and subsequent 28d recovery, PNN staining returns to normal levels (Figure 8M-N), indicating that END repopulating cells share similar PNN- regulating capacities as homeostatic microglia.

Accumulating evidence indicates that WM and developing microglia, contribute to myelinogenesis/oligodendrocyte progenitor maintenance^64–66^; thus, we next explored the effects of filling the brain with repopulating cells on WM and WM-associated cells. However, unlike during development in which these cells participate in oligo/myelinogenesis, in 28d recovery mice we observe no changes in Olig2, a marker for cells of oligodendrocyte lineage (Figure S5A, D), PDGFRα, a marker for oligodendrocyte progenitor cells (Figure S5B, E), or MBP, a marker for myelin (Figure S5C, F-G). Together, these data provide evidence that although END repopulating cells maintain an altered phenotypic profile, they can still fill the empty microglial niche with few functional consequences.

### Filling the brain with END repopulating cells reduces amyloid load in specific brain regions in the 5xFAD mouse model of Alzheimer’s disease

Given our ability to fill the brain with a subset of microglia that originate from SVZ/WM brain regions, we next explored whether these cells play distinct roles in disease pathogenesis. An early postnatal phagocytic subset of microglia located in WM (termed PAMs) has been identified, which shares a gene signature with DAMs^20^. DAMs upregulate genes involved in phagocytosis and, in humans, WM microglia express higher levels of phagocytosis-related proteins, indicating the higher phagocytic potential of these cells compared to their grey matter counterparts^15, 67^. Moreover, reports indicate that amyloid plaque pathology is exceedingly rare in WM areas in the AD brain compared to grey matter areas^68^. Since our SVZ/WM-derived END repopulating cells display DAM signatures even after 28 days of residency in grey matter areas of the brain, we were interested in exploring whether we could replace endogenous microglia with these repopulating myeloid cells in the 5xFAD mouse model prior to plaque development, and whether this replacement would impact plaque formation.

5xFAD transgenic mice harbor five familial AD-associated mutations, which result in the formation of amyloid plaques, as well as gliosis and neuronal loss, disease hallmarks of AD. WT and 5xFAD mice at 1.5 months (an age that precedes amyloid deposition) were treated for 14d with PLX3397 (600 ppm) and then allowed to recover on control diet for 3 months (Figure 9A,C). In a recent study, we showed that repopulation of microglia following 10 wk PLX5622 1200 ppm administration (which does not achieve END repopulation) in 1.5 mo 5xFAD mice restores plaque pathology following CSF1R inhibition, indicating that PND microglia seed plaques and do not differentially modulate plaques compared to endogenous microglia^69^. To show that 14d PLX3397 (600 ppm) achieves END repopulation in 5xFAD mice, mice were examined immediately following microglial depletion and during the early stages of END repopulation to confirm that elimination and subsequent END repopulation occurs in 5xFAD mice (Figure 9B). Following 3 mo of recovery, END repopulating myeloid cells still exhibit phenotypic alterations in WT mice, including in IBA1^+^ expression intensity and process diameter (Figure 9D-I). Microglia in 5xFAD mouse brains present elevated cell numbers, IBA1^+^ expression intensity, process diameter, and reduced process length relative to WT microglia; reflective of the activated and amoeboid cell phenotypes reported in AD^70, 71^. END repopulation in 5xFAD mice results in the return of microglial cell densities and process length size to WT levels (Figure 9E,I), indicating that repopulation can partially reverse some microglial morphological alterations induced by AD. Analysis of amyloid plaques, as seen by ThioflavinS+ staining, reveals a significant reduction in amyloid load in the cortex and amygdala (Figure 9D, J-K, N-O). No significant changes were observed in the thalamus (Figure 9M-N). Interestingly, we observe that there are significantly fewer IBA1^+^ cells surrounding plaques in repopulated 5xFAD mice compared to 5xFAD controls (Figure 9P). Together, these findings indicate that myeloid cells play critical roles in plaque formation, maintenance, and pathogenesis, and that END repopulating myeloid cells impair the build-up of plaque pathology compared to endogenous microglia.

**Figure 9.**
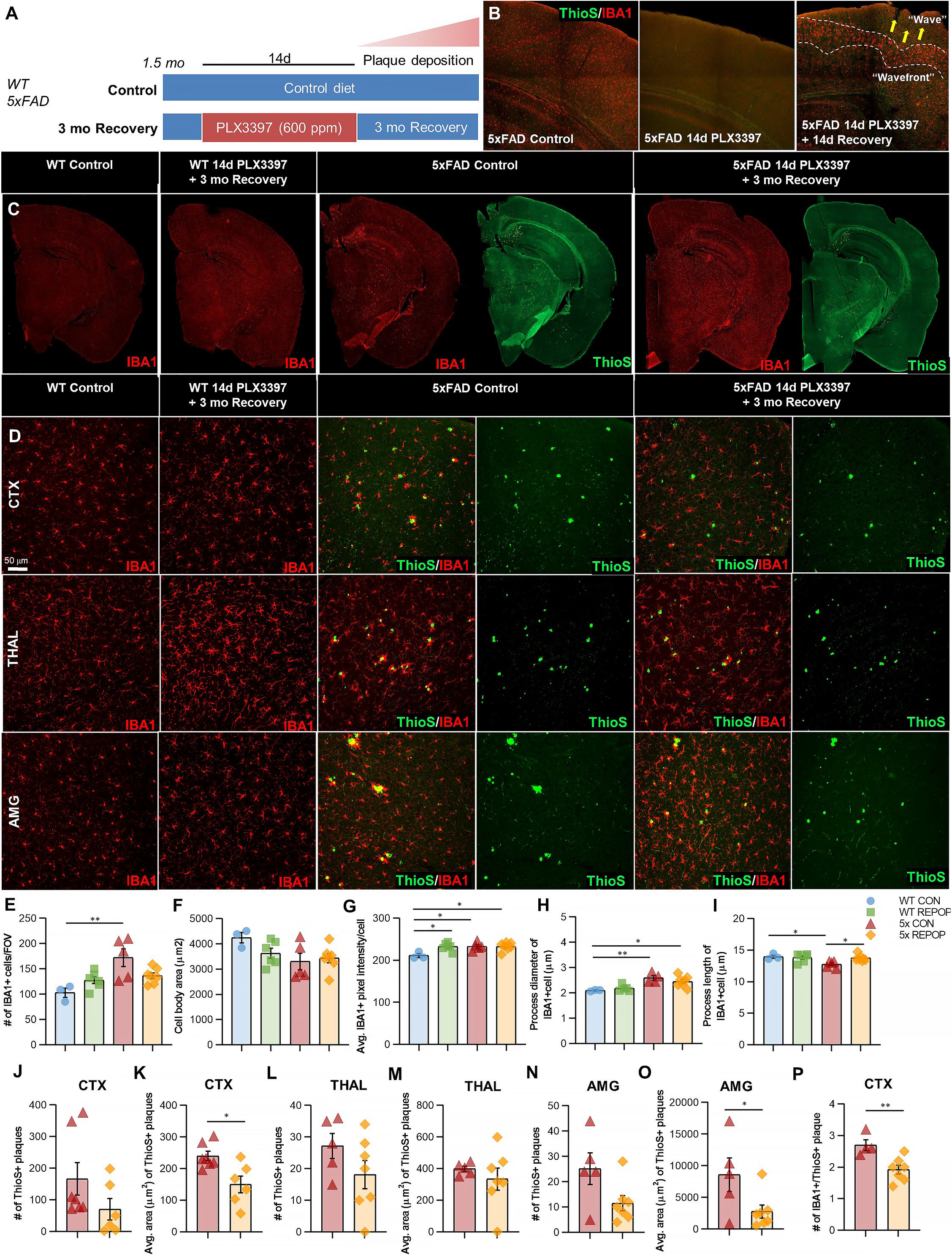
**Filling the brain with alternative repopulating cells reduces amyloid load in specific brain regions in the 5xFAD mouse model of Alzheimer’s disease.** (A) Experimental paradigm and schematic depicting dose and duration of PLX3397 (600 ppm) treatment and subsequent inhibitor withdrawal in WT and 5xFAD mice. 1.5-month-old WT or 5xFAD mice (an age that precedes plaque deposition) were treated with 600 ppm of PLX3397 for 14d, and then placed on control diet for 3 mo allowing for alternative repopulation. (B) Representative immunofluorescence images of microglia (IBA1, red) and ThioS^+^ (amyloid plaques, green) in the cortex of control, 14d PLX3397 (600 ppm), and 14d recovery 5xFAD mice. (C) Representative immunofluorescence whole brain images of microglia (IBA1, red) and ThioS^+^ (amyloid plaques, green) in control and 3mo recovery WT and 5xFAD mice. (D) Representative 20x immunofluorescence images of microglia (IBA1, red) and ThioS^+^ (amyloid plaques, green) in the cortex, thalamus, and amygdala in control and 3mo recovery WT and 5xFAD mice. (E-I) Quantification of IBA1^+^ cells and morphology: IBA1^+^ cells per field of view (E), cell body area (F), average IBA1^+^ pixel intensity (G), average process/filament diameter (H), and average process/filament length (I). (J-O) Quantification of the number of ThioS^+^ plaques (J, L, N) and average area of ThioS+ plaque coverage (K, M, O) in the cortex (J-K), thalamus (L-M), and amygdala (N-O). (P) Quantification of the number of IBA1+ microglial cells per ThioS+ amyloid plaque. Data are represented as mean ± SEM (n=3-6). * p < 0.05, ** p < 0.01. CTX, cortex; THAL, thalamus; AMG, amygdala.

## Discussion

As central players in CNS homeostasis, defense, and disease, intense focus has recently been placed on microglia and our understanding of their cell origins, function and dynamics. For decades the identity and ontogeny of microglial precursors remained controversial; the scientific community debated whether microglia derive from embryonic progenitors or blood-derived monocytes^72, 73^. It is now well-established that microglia arise from yolk sac-derived erythromyeloid progenitors^4, 5^.

We previously reported that adult microglia are dependent on CSF1R signaling for their survival and identified several CSF1Ri that eliminate microglia for extended periods of time without peripheral cell infiltration^23^. Following CSF1Ri-dependent microglial depletion, we and others have shown that subsequent withdrawal of CSF1Ri from the microglial-depleted brain results in rapid microglial repopulation derived from surviving microglia^23, 25, 29, 32, 59, 74^. The resultant tissue is reconstituted within 14-21d in a homogenous, tile-like fashion with the replacing microglia fully resembling the original tissue^23, 25, 31, 32^. Due to the rapid proliferation of these surviving microglia, exploration into the contribution of specific myeloid cell subtypes to the adult CNS has proven difficult, and so we set out to develop a paradigm without notable surviving microglia.

Previous attempts to achieve an empty microglial niche have fallen short, reporting ∼95% or less microglial depletion efficiency^41^. Here, we utilize sustained high-dose CSF1Ri administration (specifically 14d PLX3397 600 ppm) and show we can obtain 99.98% microglial depletion. In doing so, we identify a CNS myeloid cell subset that repopulates the brain parenchyma from SVZ/WM areas, without contributions from the periphery. We describe this form of repopulation as empty niche-dependent (END) repopulation due to its unprecedented level of depletion efficiency and distinct characteristics from partial microglial niche-dependent (PND) repopulation. Unlike PND repopulation paradigms, in which surviving microglia proliferate in clusters to give rise to new microglia^26^ or more uniformly throughout the brain^23, 25^, END repopulation dynamics involve specific spatiotemporal patterns and a dynamic migratory wave of proliferating cells. In addition, previous studies report a major contribution of peripheral bone marrow-derived myeloid cells to the repopulating cell population following an empty microglial niche or the persistent loss of microglia (in which microglia cannot repopulate the niche)^41, 42^. Our data refutes these findings. These models rely on tamoxifen administration thus, it could be possible that BM-derived myeloid cell engraftment in the CNS results from an experimental or technical caveat related to toxin administration rather than the presence of an empty microglial niche. In line with this, a recent study has reported that tamoxifen expands macrophage populations and should be reconsidered as a neutral agonist in myeloid cell lineage studies^75^. Recent scrutiny has also been placed on the use of CSF1Ri, implicating long-term changes in bone-marrow derived macrophages^45^. In this study, we show that high dose and sustained CSF1Ri can result in alterations to monocyte precursor populations, however, these changes do not translate in significant changes to peripheral monocyte populations, which we again show do not contribute to CNS myeloid cell repopulation in the absence of toxin, irradiation or injury.

END repopulating cells initially appear in specific neuroanatomical niches (first in the SVZ – the site of where surviving CSF1Ri resistant SVZ/WM myeloid cells reside) and spread throughout the brain in a distinct pattern: via WM tracts to the caudoputamen, optical tract, internal capsule, cerebral peduncle, and finally to cortical areas. The caudoputamen is closely associated with the lateral ventricle, SVZ and WM tracts in which myeloid cells initially appear and migrate, and we believe this spatial association plays a large role in why certain parenchymal areas see more appreciable repopulating cell deposition. Notably, this distribution of initially repopulating cells in ventricular regions and subsequent migration pattern exhibit strong anatomical parallels to microglial colonization and distribution of the embryonic and postnatal brain, in which microglia enter the brain via ventricular routes and remain restricted in WM zones before migrating out to the rest of the brain, with the cortical plate being one of the last areas of colonization^2, 7, 10, 12, 76, 77^. Del Rio-Hortega observed the accumulation of microglia along ventricles and white matter areas in the developing brain within the first postnatal week^72, 78^. In 1939, Kershman used the term “fountain of microglia” referring to “hot spots” of activated microglia during human embryogenesis. This observation has been confirmed by several studies over the decades, and recent publications have identified a distinct microglia population that appears postnatally in myelinated regions^65, 79, 80^. Thus, it could also be possible that END repopulating cells and their migratory dynamics could represent an additional wave of myeloid cells that occurs postnatally. However, it should be noted that we do not believe END repopulating cells represent a microglial precursor. Most importantly, our findings highlight important anatomical structures that facilitate microglial/myeloid cell migration in an empty microglial niche, which play an essential role in development, but also appear intact in the adult brain.

Utilizing a microglial depletion and repopulation paradigm that successfully achieves an empty microglial niche, we identified a unique subset of myeloid cells in the SVZ/WM area that serves as the source for END repopulating cells. Notably, our novel paradigm results in the replacement of microglia with *Cx3cr1*^+^ myeloid cells originating from the SVZ and associated WM areas, allowing us to study the biology of these cells, and how they adapt to extrinsic environmental cues from grey matter, rather than the WM areas they are normally restricted. These SVZ/WM myeloid cells are resistant to CSF1Ri, which we believe owes to unique properties of the SVZ environment, as once END repopulating cells (deriving from SVZ/WM myeloid cells) take up residence in other parenchymal areas, they once again are susceptible to CSF1Ri. END repopulating cells maintain altered phenotypes and responses to LPS, even after extended periods of time, suggesting that these SVZ-derived cells are highly distinct from other microglia. In line with identifying distinct properties of myeloid cells in the SVZ/WM, previous studies have reported the unique heterogeneity of this microglial subset compared to subsets in other brain regions^81, 82^. Similar to the morphological and molecular findings in END repopulating cells, SVZ microglia are less branched (or less ramified), express surface markers that are commonly associated with an alternatively active phenotype (i.e. expressing high levels of anti-inflammatory cytokines IL-4 and IL-10), and lower expression levels of P2RY12^33^. Interestingly, the existence of *Cx3cr1*^+^ IBA1^-^ cells in the SVZ and RMS has been reported^33^. In humans, microglia in the SVZ exhibit a more activated phenotype that was distinct from all other brain regions, and show higher expression of CD45, CD64, CD68, CX3CR1, EMR1, HLA-DR, indicative of microglial activation, and proliferation markers like Ki67 compared to other subsets^83^. Moreover, WM microglia have been reported to display unique properties during postnatal stages, including an amoeboid morphology and enriched expression of genes related to microglial priming, phagocytosis, and migration^84^. Thus, it appears that END repopulating myeloid cells fit many of the previously reported characteristics of SVZ/WM microglia.

Our results show that SVZ/WM microglia have the capacity to leave their brain region of residence, given certain cues and circumstances, and that these cells potentially play roles in health and disease. Once established in the brain, the gene expression profile of the END repopulating cell shows enhanced enrichment for DAM genes (e.g. *Clec7a*, *Axl*, *Ly9*, *Apoe*, *Itgax)*. DAMs have been identified as a unique set of microglia that restrict Alzheimer’s disease (AD) progression^15^. It is established that microglia are critical for the pathogenesis of AD; their depletion from the AD brain prevents the parenchymal accumulation of plaques and instead promotes Aβ deposition in the vasculature^69^. In addition, studies indicate that these cells play an essential role in synaptic and neuronal loss^85–87^, further driving disease progression. Although abnormalities in WM are currently under investigation, extensive evidence has shown that plaque deposition occurs almost exclusively in grey matter areas in the human brain^68^, despite elevated Aβ levels in WM^88^. Here, we demonstrate that replacing microglia with SVZ/WM-derived myeloid cells impedes plaque development in specific grey matter brain regions. Previous studies have shown that neither PND microglia repopulation nor replacement with monocytes has a significant impact on plaque pathogenesis^69, 89–92^, highlighting the significance of these SVZ/WM-derived myeloid cells in disease. An elegant study by Füger et al. (2017) indicates that the accumulation of plaque-associated microglia derives from local microglial proliferation^93^, thus we postulate that it is unlikely that plaque-associated microglia derive or travel from the WM tracts in untreated AD brains. Together, our results show how a specific myeloid cell population can influence pathological development and lend insight into the protective capabilities of 1) SVZ/WM myeloid cells, and 2) DAM phenotypes, against plaque formation.

In conclusion, this study unveils the presence of a myeloid cell subtype originating from an important neurogenic niche, the SVZ - and associated WM areas, with tremendous chemotaxis potential and the ability to fully reconstitute the microglial-depleted brain. These cells exhibit distinct properties compared to homeostatic microglia, sharing transcriptional profiles with DAMs and offering an ability to lower amyloid load in a mouse model of AD. Together, these results not only highlight the complexity and diversity of myeloid cells in the adult brain, but we establish a model system that provides new insight on myeloid cell homeostasis and dynamics in the brain.

## Materials and Methods

Detailed methods are provided in the Supplemental Information portion of this paper.

## Acknowledgments

We thank Edna Hingco and Ayer Darling Jue for their excellent technical assistance, and Brian L. West and Andrey Rymar at Plexxikon, Inc. for providing and formulating CSF1Ri chow and pharmacokinetics analysis. We are grateful to Claudia I. Czimczik for proofreading the manuscript. This work was supported by the National Institutes of Health (NIH) under awards: R01NS083801 (NINDS), R01AG056768 (NIA), and P50AG016573 (NIA) to K.N.G; F31NS108611 (NINDS) to J.D.C and T32NS082174 to Y.G. L.A.H. was supported by the Alzheimer’s Association Research Fellowship (AARF-16-442762).

## Author Contributions

L.A.H. developed experimental protocols, designed, performed, and analyzed experiments and wrote the manuscript. A.R.N. designed, performed and analyzed experiments. Y.G. designed, performed, and analyzed experiments. N.S. and S.J. performed bioinformatic analysis. J.D.C. designed, performed, and analyzed experiments. D.X.F.V. prepped, imaged, and analyzed the two-photon data as well as performed the skull removal surgery. S.E.R. developed experimental protocols and performed experiments. S.J.K., M.R.E., and C.M.H. performed experiments. A.J.A., S.P.G., A.M., and M.A.I. contributed to project design. K.N.G directed the project, designed the experiments, interpreted the results and wrote the manuscript.

## Supplementary Information

### Supplemental Information - Materials and Methods

#### Compounds

PLX3397 was provided by Plexxikon Inc. (Berkeley, CA) and formulated in AIN-76A standard chow by Research Diets Inc. at the doses indicated in the text. PLX3397 was provided in chow at 600 ppm.

#### Mice

All mice were obtained from The Jackson Laboratory (Bar Harbor, ME) unless otherwise indicated. *Cx3cr1^CreERT2^* (020940), *Ascl1^CreERT2^* (012882), and *Nestin^CreERT2^* (016261) mice were bred to R26-YFP (006148) reporter mice. Cx3cr1^GFP/GFP^Ccr2^RFP/RFP^ (032127) mice were bred to C57BL/6 to obtain Cx3cr1^GFP/+^Ccr2^RFP/+^ mice. H2K-*BCL*-2 transgenic mice were gifted from Irving Weissman. Ccl2 KO (004434) mice were obtained from The Jackson Laboratory. For transplant studies, bone marrow cells were isolated from CAG-EGFP mice (006567). All other mice were male C57BL/6 (000664) mice. 5xFAD mice were obtained from MMRRC-JAX (34848) and have been previously described in detail^1^. For 5xFAD genotyping, the primer sequences used were PS1 Forward 5′ - AAT AGA GAA CGG CAG GAG CA – 3′ and PS1 Reverse 5′ - GCC ATG AGG GCA CTA ATC AT – 3′. Animals were housed with open access to food and water under 12h/12h light-dark cycles. All mice were aged to 1.5 months unless otherwise indicated.

#### Animal Treatments

All rodent experiments were performed in accordance with animal protocols approved (AUP-17-179) by the Institutional Animal Care and Use Committee at the University of California, Irvine (UCI). *Microglial depletion:* Mice were administered ad libitum with PLX3397 at a dosage of 600 ppm (to eliminate microglia) or vehicle (control) for 14 days. To stimulate repopulation, PLX3397 was withdrawn and replaced with vehicle. *LPS treatment:* Lipopolysaccharide (LPS; *Escherichia coli* 0111:B4; L4130, Sigma-Aldrich, St. Louis, MO) was dissolved in phosphate-buffered saline (PBS) and administered intraperitoneally (IP) at a dose of 0.33 mg/kg animal body weight either 6 or 24 hours prior to sacrifice. *BrdU labeling:* Bromodeoxyuridine (BrdU; 000103, Thermo Fisher Scientific, Waltham, MA) was administered at a dose of 1 ml/100 g body weight (per manufacturer’s instructions) for four consecutive days. Mice were sacrificed 24 hours following last BrdU injection. *Tamoxifen treatment:* Tamoxifen (10540-29-1, Sigma-Aldrich) was suspended in corn oil for 60 min at 50°C. To obtain efficient conversion of *loxP* alleles a dose of 5 mg tamoxifen/ 25 g animal body weight was delivered orally over five consecutive days. Animals were injected with tamoxifen immediately following PLX3397 inhibition to track the lineage of the repopulating cells (for all lines except *Cx3cr^CreERT2^*, in which TAM was administered 21 days prior to PLX3397 treatment). *In vivo neutralization of CCL12:* 100 ug of polyclonal goat anti- CCL12/MCP-5 (AF428, R&D Systems, Minneapolis, MN) or goat IgG in 0.5 ml of sterile HBSS was administered per mouse via i.p. injection at day 1, day 3, day 5 and day 6 (25 ug x 4) recovery (i.e. days post PLX3397 withdrawal). *Bone marrow transplant:* C57BL/6 mice were anesthetized with isoflurane and then irradiated with 1000 cGy (head-shielded) and reconstituted via retroorbital injection with 2 x 10^6^ whole BM cells from CAG-EGFP mice. Blood was measured 4, 8, and 12 weeks post transplantation to track granulocyte chimerism. At 12 weeks post transplantation, the mice were euthanized and BM was harvested and analyzed by flow cytometry for HSC chimerism. This established an average percent chimerism of >40% in HS irradiated mice. *Tissue collection:* Following treatments, adult mice were sacrificed via carbon dioxide inhalation and perfused transcardially with 1X PBS. For mouse pups (below the age of postnatal day P9), animals were fully sedated using ice and then decapitated. Brains were extracted and dissected down the midline, with one half flash-frozen for subsequent RNA and protein analyses, and the other half drop-fixed in 4% paraformaldehyde. Fixed brains were cryopreserved in PBS + 0.05% sodium azide + 30% sucrose, frozen, and sectioned at 40 μm on a Leica SM2000 R sliding microtome for subsequent immunohistochemical analyses. *Subventricular zone microdissection and isolation:* Extracted brains were immersed in ice-cold HBSS (14025092, Thermo Fisher Scientific) and cut in half along the sagittal axis. Following removal of the septum, a thin layer of the rostral and lateral walls of the lateral ventricles were extracted from each hemisphere with a microsurgical stab knife (52-1501, Unique Technologies, Mohnton, PA) and immediately frozen in RNA isolation buffer solution.

#### Histology and confocal microscopy

Fluorescent immunolabeling followed a standard indirect technique as described previously^2^. Brain sections were stained with antibodies against: IBA1 (1:1000; 019-19741, Wako Chemicals, Richmond, VA; and ab5076, Abcam, Cambridge, UK), CD11b (1:50, MCA711, Bio-Rad Laboratories, Hercules, CA), P2RY12 (1:200, HPA014518, Sigma-Aldrich), TMEM119 (1:200, ab209064, Abcam), AXL (1:100, AF854, R&D Systems), Dectin-1 (also known as CLEC7A; 1:30, mabg-mdect, Invivogen, San Diego, CA), Ki67 (1:200, ab16667, Abcam), NESTIN (1:200, ab6142, Abcam), MASH1 (1:200, 556604, BD Biosciences, San Jose, CA), TIE2 (1:100, ab24859, Abcam), GFAP (1:3000, ab4674, Abcam), DCX (1:200, sc-8066, Santa Cruz, Dallas, TX), OLIG2 (1:200, ab109186, Abcam), SOX2 (1:200, AF2018, R&D Systems), YFP/GFP (1:200, ab6556, Abcam), GFP (1:200, ab13970, Abcam), PU.1 (1:200, 22585, Cell Signaling Technology, Danvers, MA), Collagen IV (1:200, ab6586, Abcam), Parvalbumin (1:200, MAB1572, Sigma-Aldrich), and WFA (1:1000, B-1355, Vector Laboratories, Burlingame, CA). For DAPI staining, mounted brain sections were cover-slipped using Fluoromount-G with DAPI (00-4959-52, Invitrogen, Carlsbad, CA). For Thioflavin-S (Thio-S) staining, sections were washed with 1X PBS (1 × 5 min), dehydrated in a graded series of ethanol (100%, 95%, 70%, 50%; 1 × 3 min each), and then incubated in 0.5% Thio-S (in 50% ethanol, T1892, Sigma-Aldrich) for 10 min. This was followed by 3 × 5 min washes with 50% ethyl alcohol and a final wash in 1X PBS (1 × 10 min). High resolution fluorescent images were obtained using a Leica TCS SPE-II confocal microscope (Leica Microsystems, Wetzlar, Germany) and LAS-X v 3.3.0 software. Images in the cortex were taken in the somatosensory cortex unless otherwise indicated. For confocal imaging, one field of view (FOV) per brain region was captured per mouse unless otherwise indicated. To capture brightfield images and whole brain stitches, automated slide scanning was performed using a Zeiss AxioScan.Z1 equipped with a Colibri camera (Zeiss, Oberkochen, Germany) and Zen AxioScan 2.3 software. Microglial morphology was determined using the filaments module in Bitplane Imaris 7.5 (Bitplane, Zurich, Switzerland), as described previously^3^. Cell quantities were determined using the spots module in Imaris. Percent coverage measurements were determined in Image J (NIH, Bethesda, MD).

#### Cranial Window Implantation

Mice were anesthetized with isoflurane (Patterson Veterinary, Greeley, CO) in O2 (2% for induction, 1-1.5% for maintenance). To provide perioperative analgesia, minimize inflammation, and prevent cerebral edema, Carprofen (10 mg/kg, s.c., Zoetis, Parsippany-Troy Hills, NJ) and Dexamethasone (4.8mg/kg, s.c. Phoenix Pharmaceuticals, St. Joseph, MO) were administered immediately following induction. Ringer’s lactate solution (0.2mL/20g/hr, s.c, Hospira, Lake Forest, IL) was given throughout the surgery to replace fluid loss. Sterile eye ointment (Rugby Laboratories, Hempstead, NY) was applied to prevent corneal drying. Surgical tools were sterilized using a hot glass bead sterilizer (Germinator 500, CellPoint Scientific, Gaithersburg, MD). Following hair removal, Povidone-iodine (Phoenix) and Lidocaine Hydrochloride Jelly (2%, Akorn, Lake Forest, IL) was used to disinfect and numb the scalp, respectively. The scalp and underlying connective tissue were removed to expose the parietal and interparietal bone. Lidocaine Hydrochloride injectable (2%, Phoenix) was used for muscle analgesia and the right temporal muscle detached from the superior temporal line. The skull was dried using ethanol (70% in DI water) and a thin layer of Vetbond Tissue Adhesive (3M, Saint Paul, MN) applied to the exposed surface. Custom-printed ABS headplates were attached using Contemporary Ortho-Jet liquid and powder (Lang Dental, Wheeling, IL) at an angle parallel to the skull. A small craniotomy (3mm diameter) was performed over the right hemisphere 2.5mm anterior and 3mm lateral lambda. Hemostatic gelfoam sponges (Pfizer, New York, NY) pre-soaked in sterile saline (CareFusion AirLife Modudose, CareFusion/BD, San Diego, CA) were used to absorb dural bleeding. Surgery was terminated if dural tears or intracerebral bleeding was observed. A 4mm glass coverslip (World Precision Instruments, Sarasota, FL) was placed over the exposed brain and its edges attached to the skull first with a thin layer of Vetbond and second with dental acrylic. Following surgery, mice recovered in their home cage over a warm heating pad until normal behavior resumed (∼15-30 minutes). Postoperative care consisted of daily Carprofen injections (10mg/kg, s.c.) for one week.

#### Two-Photon Imaging

Fluorescence was gathered with a resonant two-photon microscope (Neurolabware, Los Angeles, CA) with 900 nm excitation light (Mai Tai HP, Spectra-Physics, Santa Clara, CA). A 20x water immersion lens (1.0 NA, Olympus, Tokyo, Japan) was used with magnification 4. Emissions were filtered using a 510/84nm and 607/70 nm BrightLine bandpass filter (Semrock, Rochester, NY). Image sequences were gathered using Scanbox acquisition software (Scanbox, Los Angeles, CA) at a depth of 200-260µm below the meninges. An electrically tunable lens was used to image 20 planes (326x325µm, 3µm z step), each sampled at 0.5Hz. Laser damage consisted of line scanning at magnification 25 for 1-10s at 800nm.

#### Quantification of Homeostatic Motility and Response to Laser Ablation

All image stacks were processed and analyzed using the image processing package FIJI, a distribution of NIH Image J software (Schindelin et al., 2012). Stacks were temporally binned by taking the sum for each pixel over 30 frames (1 min.). Homeostatic motility was quantified manually by measuring the difference of visibly moving processes over 2 minutes. For each mouse, 10 microglia were chosen at random and the first 5 extension or retraction observed were recorded for a total of 50 observations per mouse. Microglia respond to laser damage by extending their processes toward the site of injury. We took advantage of increasing fluorescence at the site of damage from infiltrating GFP-positive processes to quantify microglia response to laser ablation. The average GFP intensity within a circle (r = 50µm) centered at site of damage at any timepoint (tx) was normalized to the intensity in that area at 1 min. post ablation (t1). To determine differences in GFP intensity within groups over time and between groups at any timepoint we used a repeated measures two-way ANOVA corrected for multiple comparison (Geisser-greenhouse correction).

#### PK Analysis

PLX3397 concentration in plasma and cerebellum were analyzed for pharmacokinetic (PK) data by Integrated Analytical Solutions, Inc.

#### FACS Analysis

Myeloid cells were extracted from whole hemispheres, isolated into single-cell suspensions and identified using fluorescence-activated cell sorting (FACS) gating for CD11b^+^CD45^int^ as previously described^2^. Cells were stained with the following surface antibodies purchased from Biolegend (San Deigo, CA) at 1:200 unless otherwise indicated: CD34-eFlour660 (1:50, 50-0341-80, eBioscience, San Diego, CA), Sca-1-AF700 (1:100, 108141), CD16/32-PE (101307), Ter119-PE/Cy5 (116209), ckit/CD117-PE/Cy7 (25-1171-81, eBioscience), CD150/SLAM-PerCP-eFlour710 (46-1502-82, eBioscience), CD11b-APC (101212), CD11b-PE (101208), Gr1-AF700 (108422), CD45-AF700 (103128), CD45-APC/Cy7 (103116), NK1.1-PE (108707), CD3-PE/Cy7 (100220), CD19-Per-Cyanine5.5 (45-0193-82, eBioscience), CD11c-APC/Cy7 (117323), Ly6C-PE (1:400, 128007), Ly6G-5.5 (127615). Samples were acquired with the BD LSRII or BD Fortessa X20, and sorted with the BD FACS Aria II.

#### RNA-sequencing and analysis

Total RNAs were extracted by using RNeasy Mini Kit (Qiagen, Hilden, Germany). RNA integrity number (RIN) was measured by Agilent 2100 Bioanalyzer (Agilent Technologies, Santa Clara, CA) and samples with RIN >= 7.0 were kept for cDNA synthesis. cDNA synthesis and amplification were performed followed by Smart-seq2^4^ standard protocol. Libraries were constructed by using the Nextera DNA Sample Preparation Kit (Illumina, San Diego, CA). Libraries were base-pair selected based on Agilent 2100 Bioanalyzer profiles and normalized as determined by KAPA Library Quantification Kit (Illumina). The libraries were sequenced using paired-end 43bp mode on Illumina NextSeq500 platform with around 10 million reads per sample. *Read alignment and expression quantification:* Pair-end RNA-seq reads were aligned using STAR v.2.5.1b with the options (--outFilterMismatchNmax 10 --outFilterMismatchNoverReadLmax 1 -- outFilterMultimapNmax 10)^5^. Rsubread was used to generate feature counts^6^. Gene expression was measured using Limma, edgeR, and org.Mm.eg.db packages with expression values normalized as RPKM^7–10^. *Differential expression analysis:* Libraries with uniquely mapping percentages higher than 80% were considered to be of good quality and kept for downstream analysis. Protein coding and long non-coding RNA genes, with expression RPKM >=1 in at least three samples, were collected for subsequent analysis. Differential expression analysis was performed by using Limma, edgeR, and org.Mm.eg.db^7–10^. Differentially-expressed genes (DEGs) were selected by using false discovery rate (FDR)<0.05. Top significant genes are displayed as a volcano plot constructed using GLimma, ggplot2, and EnhancedVolcano (FDR < 0.05, LogFC >1)^11^. PCA plots were generated using plot3D^12^. *Gene ontology and pathway analysis:* DEGs were analyzed for Gene ontology (GO) enrichment by clusterProfiler using a hypergeometric test with corrected p-values < 0.05^13^. These results were then plotted with GOplot. maSigPro package was used to identify genes that show different gene expression profiles over time^14^. Heatmaps were generated by mapping RPKM values to genes identified in maSigPro and then constructed using gplots. Normalized (min max normalization for each individual gene) log2-transformed expression values are displayed as a heatmap with hierarchical clustering utilizing gplots. maSigPro-selected gene clusters were identified and enriched for GO using clusterProfiler. GOplot was used to correlate genes and important pathways.

#### Behavioral and Cognitive Analysis

Mouse behavior, motor function, and cognition was evaluated using the following tasks: elevated plus maze, open field, novel place/novel object, sociability test, and spontaneous alternation Y-maze in the order listed, and as previously described unless otherwise indicated^2, 3, 15, 16^. Testing was conducted at 28 days recovery (i.e. after CSF1R inhibitor removal and microglial repopulation). *Sociability test:* The Crawley’s or Three-Chamber Sociability test assesses general sociability, or time spent with another rodent. In brief, animals were placed in a Three-chamber Sociability Test box (19 cm x 45 cm) with two dividing walls made of clear Plexiglas allowing free access to each chamber. During habituation, the subject mouse is placed in the middle chamber for 5 min for adaption. During testing (24 hr after habituation), a stranger mouse (inside a wire containment cup) is placed in one of the side chambers, and the subject mouse was placed in the center chamber and allowed to access and explore all three chambers for 10 min. The placement of the stranger mouse in the left and right chambers is systemically altered between trials. The duration of time spent in each chamber, velocity, and distance traveled was measured. *Spontaneous Alternation Y-Maze:* For this task, mice were placed in a Y-maze (35.2 cm arm length x 5 cm width x 20 cm sidewall height). Each animal was allowed to freely explore the arena for 8 min. Distinct intra-maze visual cues were positioned at the end of each arm for spatial orientation. Spontaneous alternation, which measures the willingness of an animal to explore new environments, was measured by the number of triads, or entry of all three arms in a consecutive sequence (i.e. ABC and not BAB). Unless otherwise indicated, behavioral readouts for all tasks were calculated from video using the EthoVision XT 14 tracking system (Noldus, Leesburg, VA).

#### Data analysis and statistics

Statistical analysis was performed with Prism Graph Pad (v.8.1.1, GraphPad Software, San Diego, CA). To compare two groups, the unpaired Student’s t-test was used. To compare multiple groups, a one-way ANOVA with Tukey’s posthoc test was performed. For all analyses, statistical significance was accepted at p < 0.05. All bar graphs are represented as means +/- SEM and significance expressed as follows: *p < 0.05, **p< 0.01, ***p < 0.001. n is given as the number of mice within each group.

## Supplemental Information – Figures and Legends

**Figure 1 - Figure Supplement 1.**
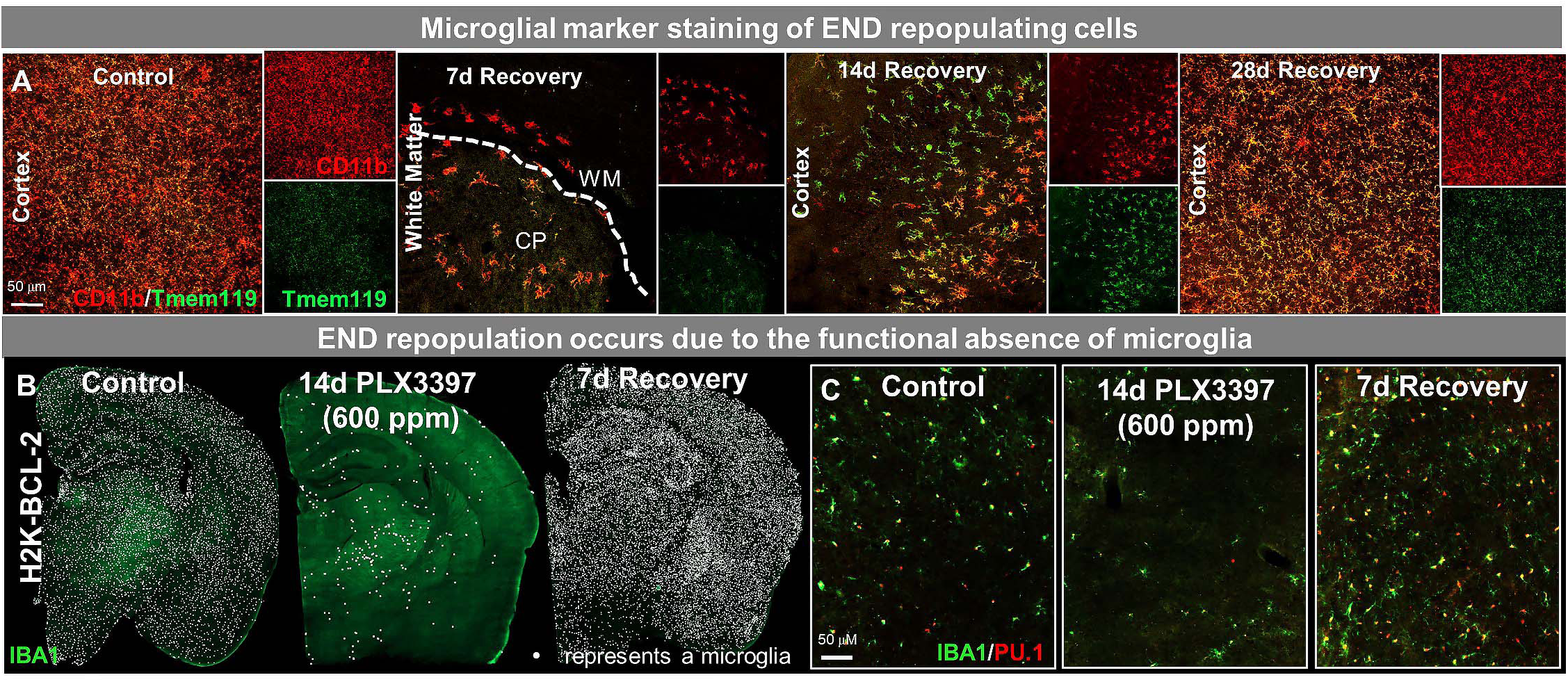
**Characterization of END repopulation: END repopulating cell staining for microglial specific marker TMEM119 and END repopulation occurs due to the functional absence of microglia in the brain parenchyma.** (A) Representative 20x images of myeloid cells (Cd11b, red) and a microglial-specific marker (P2RY12, green). (B-C) H2K-BCL2 mice were treated with PLX3397 for 14d (600 ppm in chow), the drug was withdrawn, and then mice were provided with 7 days to recover, allowing for repopulation. (B) Representative whole brain images of IBA1^+^ (green) cells in these animals, showing incomplete microglial depletion leads to remaining microglia-derived repopulation. (C) Higher resolution images of IBA1+ (green) and PU.1+ (red), a myeloid cell marker, cells.

**Figure 4 – Figure Supplement 2.**
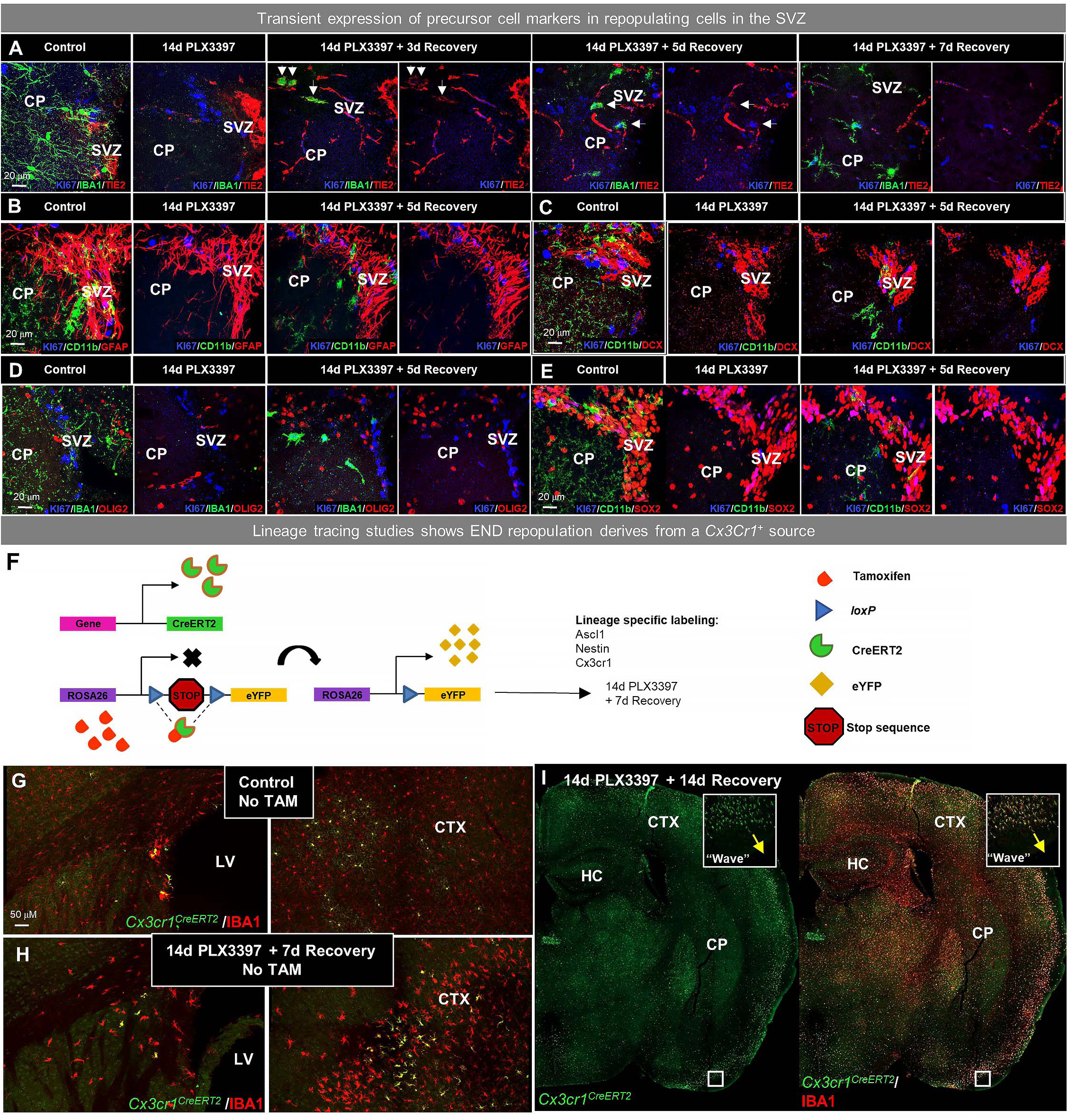
**Early END repopulating myeloid cells transiently express some precursor cell markers and Cre-lineage tracing analysis**. (A-E) Representative 63x immunofluorescence images of proliferating (Ki67^+^, blue) myeloid cells (CD11b^+^ or IBA1^+^, green) staining for positive for common cell lineage/precursor cell markers: TIE2 (red, A), and negative for: GFAP (red, B), doublecortin (DCX, red, C), OLIG2 (red, D), and SOX2 (red, E) in the SVZ of control 14d PLX3397, 3d recovery, 5d recovery, and 7d recovery mice. (F-I) CreER-directed lineage-specific labeling. (F) In these mouse lines, tamoxifen-inducible Cre-recombinase is expressed under control of the promoter of interest. When activated by tamoxifen, the CreER fusion protein translocates to the nucleus allowing transient recombination to occur and, when crossed to a YFP reporter, visualization of induced expression via eYFP. In this study, mice received 14d PLX3397 treatment 21d following last tamoxifen injection. (G-H) Representative images of *Cx3cr1^CreERT2^*-YFP^+^ and IBA1^+^ cells in control (G) and 7d recovery (H) mice treated without tamoxifen. (I) Representative whole brain images of *Cx3cr1^CreERT2^*-YFP^+^ (green) and IBA1^+^ (red) cells in 14d recovery mice, highlighting the “wave.” CTX, cortex; CP, caudoputamen; HC, hippocampus; LV, lateral ventricle; SVZ, subventricular zone.

**Figure 6 – Figure Supplement 3.**
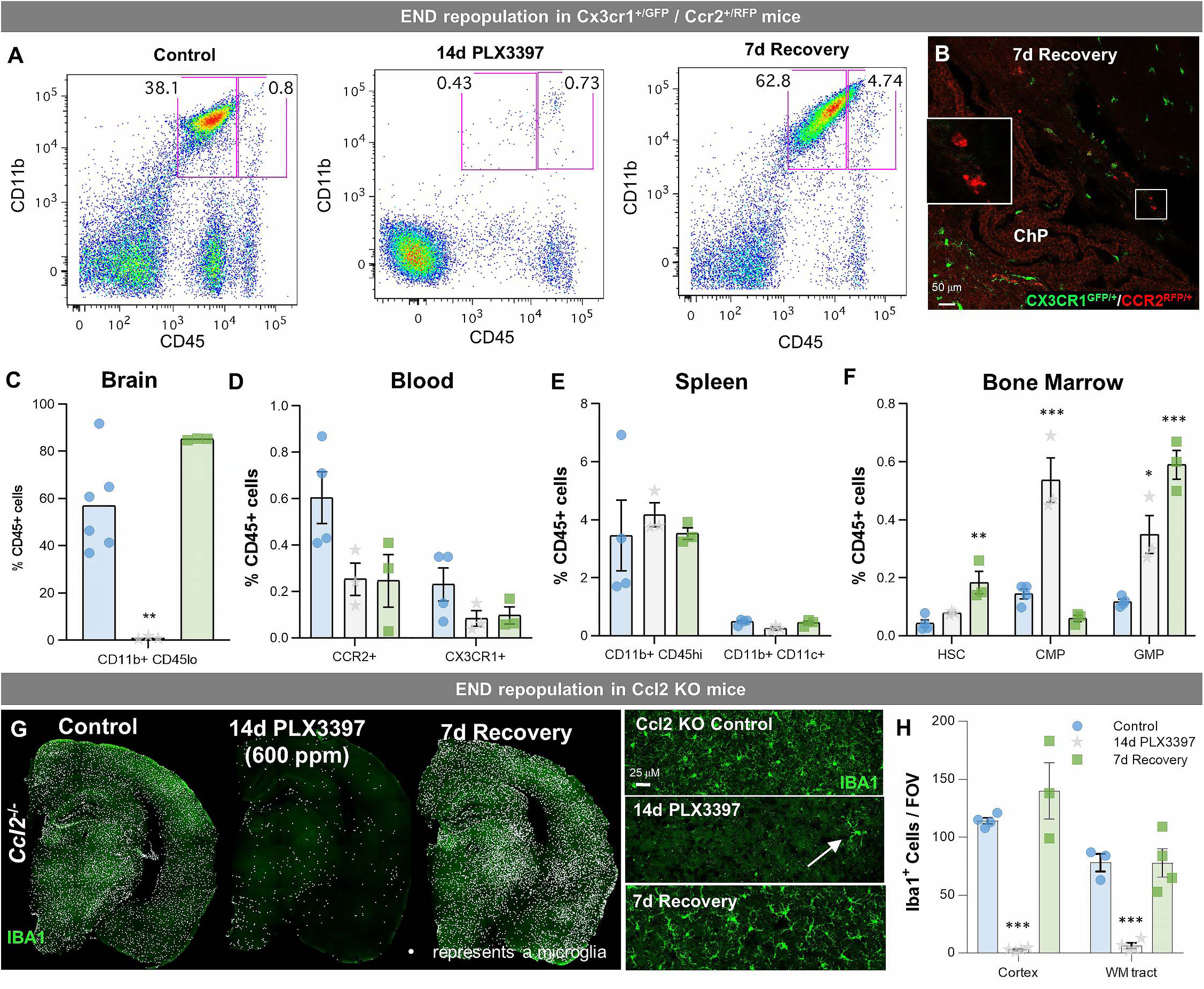
**Evaluation of peripheral cell dynamics and CCL2 signaling in END repopulation**. (A-F) Sustained microglial depletion and END repopulation in Cx3cr1^GFP/+^CCR2^RFP/+^mice. (A) Flow cytometry gating strategy for identifying microglial (CD11b^+^CD45^lo^) cells in the brain. (B) Representative immunofluorescence image of CX3CR1^+^ (green) and CCR2^+^ (red) cells in 7d recovery mice. END repopulation is apparent and no CCR2+ cells are present in the brain parenchyma. CCR2^+^ cells are only found in the choroid plexus (see insert), indicating that repopulating cells do not derive from CCR2^+^ (a common marker for peripheral myeloid cells or monocytes) cells. (C-F) Quantification of the impact of treatment on microglia in the brain (C) and distinct myeloid or myeloid precursor cell subsets in the blood (D), spleen (E), and bone marrow (F). (G-H) Ccl2 KO mice were treated with PLX3397 for 14d (600 ppm in chow), the drug was withdrawn, and then mice were provided with 7 days to recover, allowing for repopulation. (G) Representative whole brain images of IBA1^+^ (green) cells in these animals, showing incomplete microglial depletion leads to PND repopulation. Inserts show higher resolution images of IBA1+ (green) cells. (H) Quantification of the average number of IBA1^+^ cells per FOV in the cortex and white matter tract of control, 14d PLX3397 (600 pm), and 7d recovery mice. Data are represented as mean ± SEM (n=3-4). *p < 0.05, **p < 0.01, ***p < 0.001. ChP, choroid plexus; HSC, hematopoietic stem cell; CMP, common myeloid progenitor; GMP, granulocyte-macrophage progenitor.

**Figure 7 – Figure Supplement 4.**
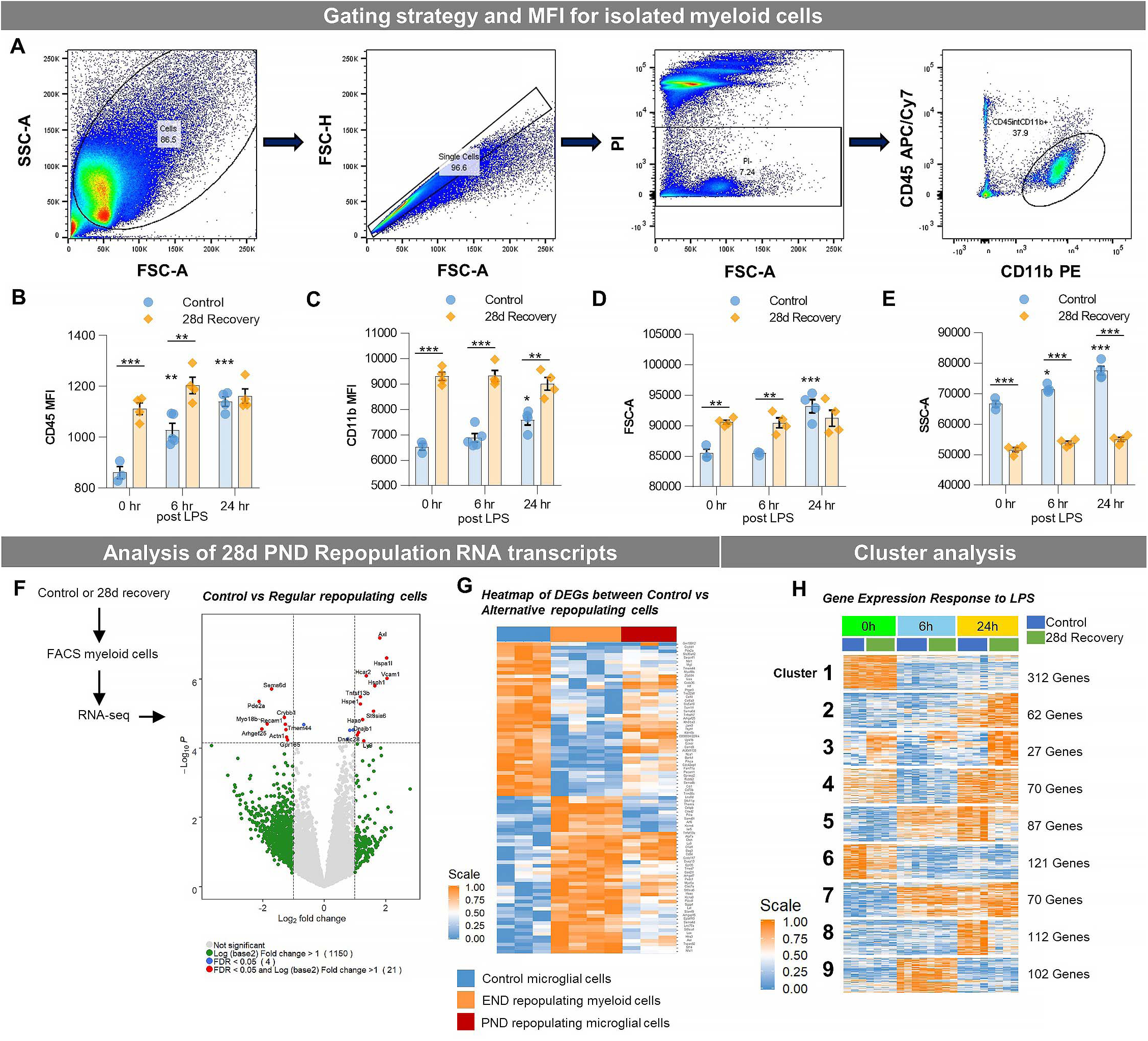
**END repopulating myeloid cells are transcriptionally distinct from homeostatic microglia.** (A) FACS gating strategy for CD11b^+^CD45^int^ myeloid cells isolated from half brains. (B-C) Quantification of CD45 and CD11b intensities of extracted myeloid cells, respectively. (D-E) Quantification of forward- and side-scatter values of extracted myeloid cells, respectively. (F-G) 2-month-old WT mice treated with vehicle (blue) or PLX3397 (600 ppm in chow) for 7 or 14 days to stimulate PND (red) and END (orange) repopulation, respectively. Microglia were isolated via FACS at 28 days recovery, RNA was extracted, and gene expression analysis was performed using RNA-seq. (F) Volcano plots displaying the fold change of genes (log2 scale) and their significance (y axis, - log10 scale) between control vs. PND repopulation microglia in 28d recovery mice. (G) Heatmap of identified differentially-expressed genes (DEGs) between control vs. 28d myeloid cells following 14d PLX3397 (600 ppm in chow; from Figure 2F), showing distinct expression patterns in control (blue), END repopulated cells (orange), and PND repopulated cells (red). (H) Heatmap of full time-series cluster analysis of control and 28d recovery cells. Provided number indicates number of genes per cluster. Data are represented as mean with individual data points (n=3).

**Figure 8 – Figure Supplement 5.**
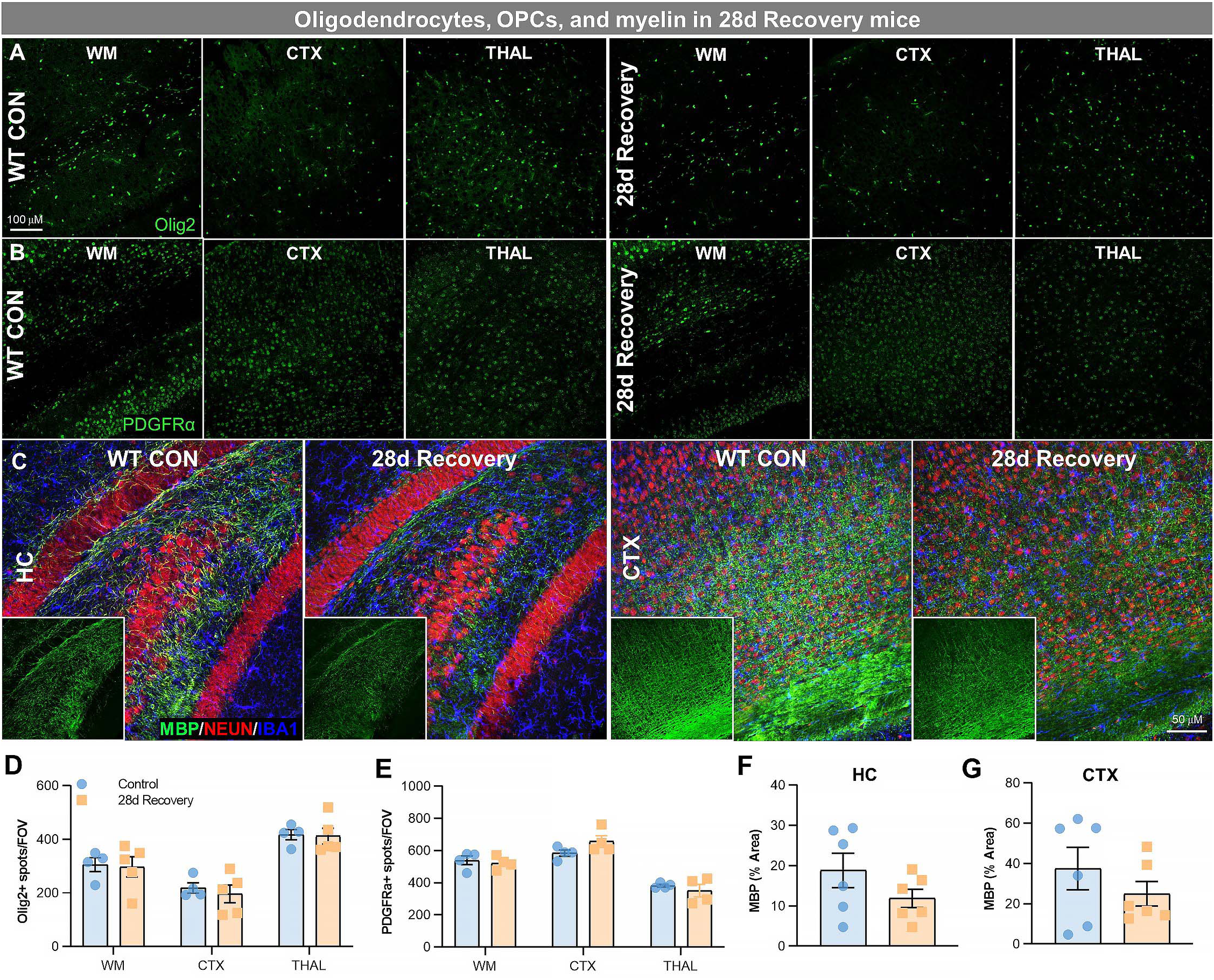
**Effects of END repopulation on white matter and cells of oligodendrocyte lineage.** (A-B) Representative 20x immunofluorescence images of Olig2^+^ (A), a marker for oligodendrocytes, and PDGFRα^+^ (B), a marker for oligodendrocyte precursor cells (OPC), cells from control and 28d recovery mice in white matter, cortical, and thalamic regions. (C) Representative 20x immunofluorescence images of myelin basic protein (MBP; green), NEUN+ cells (red, to show brain structure), and IBA1^+^ cells (blue) from control and 28d recovery mice in the hippocampus and cortex. (D) Quantification of (A). (E) Quantification of (B). (F-G) Quantification of (C). Data are represented as mean ± SEM (n=4-6). *p < 0.05, **p < 0.01. CTX, cortex; HC, hippocampus; THAL, thalamus.

**Supplemental Video 1. Cx3cr1-GFP+ myeloid cell response in control mice after laser ablation.** Representative video of Cx3cr1-GFP^+^ myeloid cell response to laser ablation (1-5 sec long) captured over a 62 min time period in control mice obtained via two-photon imaging. Each frame captures 30 sec of elapsed time.

**Supplemental Video 2. Cx3cr1-GFP+ myeloid cell response in 35d recovery mice after laser ablation.** Representative video of Cx3cr1-GFP^+^ myeloid cell response to laser ablation (1-5 sec long) captured over a 62 min time period in 35d recovery mice obtained via two-photon imaging. Each frame captures 30 sec of elapsed time.

